# IFITM proteins promote SARS-CoV-2 infection and are targets for virus inhibition

**DOI:** 10.1101/2020.08.18.255935

**Authors:** Caterina Prelli Bozzo, Rayhane Nchioua, Meta Volcic, Jana Krüger, Sandra Heller, Christina M. Stürzel, Dorota Kmiec, Carina Conzelmann, Janis Müller, Fabian Zech, Desiree Schütz, Lennart Koepke, Elisabeth Braun, Rüdiger Groß, Lukas Wettstein, Tatjana Weil, Johanna Weiß, Daniel Sauter, Jan Münch, Federica Diofano, Christine Goffinet, Alberto Catanese, Michael Schön, Tobias Böckers, Steffen Stenger, Kei Sato, Steffen Just, Alexander Kleger, Konstantin M.J. Sparrer, Frank Kirchhoff

## Abstract

Interferon-induced transmembrane proteins (IFITMs 1, 2 and 3) are thought to restrict numerous viral pathogens including severe acute respiratory syndrome coronaviruses (SARS-CoVs). However, most evidence comes from single-round pseudovirus infection studies of cells that overexpress IFITMs. Here, we verified that artificial overexpression of IFITMs blocks SARS-CoV-2 infection. Strikingly, however, endogenous IFITM expression was essential for efficient infection of genuine SARS-CoV-2 in human lung cells. Our results indicate that the SARS-CoV-2 Spike protein interacts with IFITMs and hijacks them for efficient viral entry. IFITM proteins were expressed and further induced by interferons in human lung, gut, heart and brain cells. Intriguingly, IFITM-derived peptides and targeting antibodies inhibited SARS-CoV-2 entry and replication in human lung cells, cardiomyocytes and gut organoids. Our results show that IFITM proteins are important cofactors for SARS-CoV-2 infection of human cell types representing in vivo targets for viral transmission, dissemination and pathogenesis and suitable targets for therapeutic approaches.

## INTRODUCTION

SARS-CoV-2 is the cause of pandemic Coronavirus disease 2019 (COVID-19). Originating from China in late 2019, the virus has infected more than 76 million people around the globe (https://coronavirus.jhu.edu/map.html). While SARS-CoV-2 spreads more efficiently than SARS-CoV and MERS-CoV, the previously emerging causative agents of severe acute respiratory syndromes (SARS), it shows a lower case-fatality rate (~2 to 5%), compared to ~10% and almost 40%, respectively^1–3^. The reasons for this efficient spread and the mechanisms underlying the development of severe COVID-19 are incompletely understood but the ability of SARS-CoV-2 to evade or counteract innate immune mechanisms may play a key role ^4^.

Here, we focused on innate immune effectors that are thought to target the first essential step of SARS-CoV-2 replication: entry into its target cells. A prominent family of interferon (IFN) stimulated genes (ISGs) known to inhibit fusion between the viral and cellular membranes are interferon-inducible transmembrane (IFITM) proteins^5,6^. The three best characterised members of the IFITM family are IFITM1, IFITM2 and IFITM3^7–10^. They contain different sorting motifs and IFITM1 is mainly localised at the plasma membrane, while IFITM2 and 3 are found inside the cell on endo-lysosomal membranes^7^. Thus, IFITM proteins may act at different sites of viral entry and it has been reported that they restrict multiple classes of enveloped viral pathogens including Influenza A viruses, Flaviviruses, Rhabdoviruses, Bunyaviruses and human immunodeficiency viruses^6,11^. The molecular mechanism(s) underlying the antiviral activity of IFITMs are not fully understood. However, recent reports suggest that they modulate membrane rigidity and curvature to prevent fusion of the viral and cellular membranes^12–14^.

It has also been reported that IFITM proteins inhibit human coronaviruses including SARS-CoV-1 and SARS-CoV-2 as well as MERS-CoV^11,15^. However, most results were obtained using Spike containing viral pseudoparticles and cell lines overexpressing the IFITM proteins and frequently also the viral ACE2 receptor. Here, we confirmed and expanded previous results showing that IFITM proteins block SARS-CoV-2 entry under such artificial experimental conditions. In striking contrast, however, endogenous IFTIM proteins were essential for efficient infection and replication of genuine SARS-CoV-2 in various types of human cells. We found that IFITM proteins are expressed in human cell types involved in virus transmission, dissemination to various organs, and development of severe COVID-19. In further support of an important role of IFITM proteins as entry cofactors of SARS-CoV-2, IFITM-derived peptides and targeting antibodies efficiently inhibited SARS-CoV-2 infection of human lung, heart and gut cells. Our unexpected finding that SARS-CoV-2 hijacks human IFITM proteins for efficient infection helps to explain the rapid spread of this pandemic viral pathogen.

## Results

### Overexpressed IFITMs block and endogenous IFITMs boost SARS-CoV-2 infection

It has been reported that overexpression of IFITM proteins prevents entry of viral particles pseudotyped with the Spike (S) proteins of SARS- and MERS-CoVs^9,11,15^. In agreement with these previous findings, we found that IFITM1, IFITM2 and (less efficiently) IFITM3 dose-dependently inhibited SARS-CoV-2 S-mediated entry of Vesicular-Stomatitis-Virus pseudoparticles (VSVpp) into transfected HEK293T cells (Fig. 1a, Extended Data Fig. 1a, b). Inhibition of SARS-CoV-2 S-mediated infection by IFITM proteins was confirmed using lentiviral pseudoparticles (LVpp, Extended Data Fig. 1c). In contrast, IFITMs did not significantly affect VSV-G-dependent entry (Extended Data Fig. 1d). To examine the impact of endogenous IFITM expression on S-mediated VSVpp infection, we performed siRNA knock-down (KD) studies in the human epithelial lung cancer cell line Calu-3, which expresses ACE2^16^ and increased levels of all three IFITM proteins upon IFN treatment (Extended Data Fig. 2a). On average, silencing of IFITM expression (Extended Data Fig. 2b) enhanced VSVpp infection mediated by SARS-CoV S proteins about 3- to 7-fold (Fig. 1b). To determine whether overexpression of IFITMs also affects genuine SARS-CoV-2 replication, we infected HEK293T cells overexpressing ACE2 alone or together with individual IFITM proteins. In agreement with the inhibitory effects on S containing VSVpp and LVpp, IFITM1 and IFITM2 prevented viral RNA production almost entirely, while IFITM3 achieved ~5-fold inhibition (Fig. 1c).

**Fig 1.**
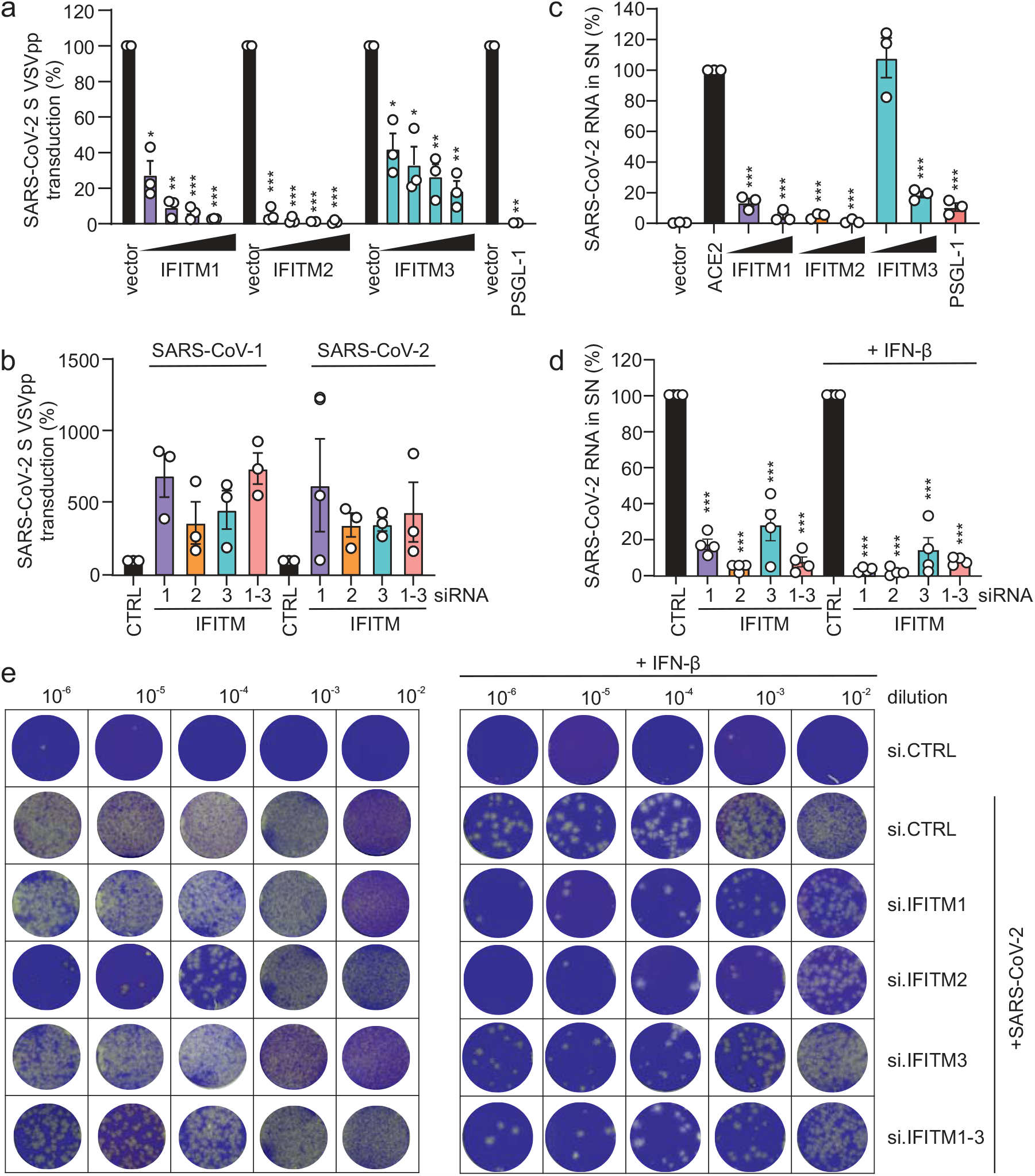
Opposing effects of IFITM proteins on SARS-CoV-2 infection. **a**, Quantification of VSV(luc)ΔG*SARS-CoV-2-S entry by measuring luciferase activity in HEK293T cells transiently expressing the indicated IFITM proteins. Bars in all panels show results of three independent experiments (mean value, ±SEM). **b**, Calu-3 cells treated with non-targeting (CTRL) or IFITM1, 2 or 3 siRNAs or a combination of the three and infected with VSV(luc)ΔG*SARS-CoV-2-S particles. **c**, Quantification of RNA containing N gene sequences by qRT-PCR in the supernatant of HEK293T cells transiently expressing ACE2 alone or together with the indicated IFITM proteins 48 h post-infection with SARS-CoV-2 (MOI 0.05). **d**, RNA containing N gene sequences levels in the supernatant of Calu-3 cells, collected 48 h post-infection with SARS-CoV-2 (MOI 0.05). Cells were transfected with control (CTRL) or IFITM1, 2 and/or 3 targeting siRNA or a combination of the three and either treated with IFN-β or left untreated as indicated. **e**, Cytopathic effects in Vero cells infected with serial dilutions of Calu-3 supernatants from Figure 1d. Cells were stained with crystal violet.

**Fig 2.**
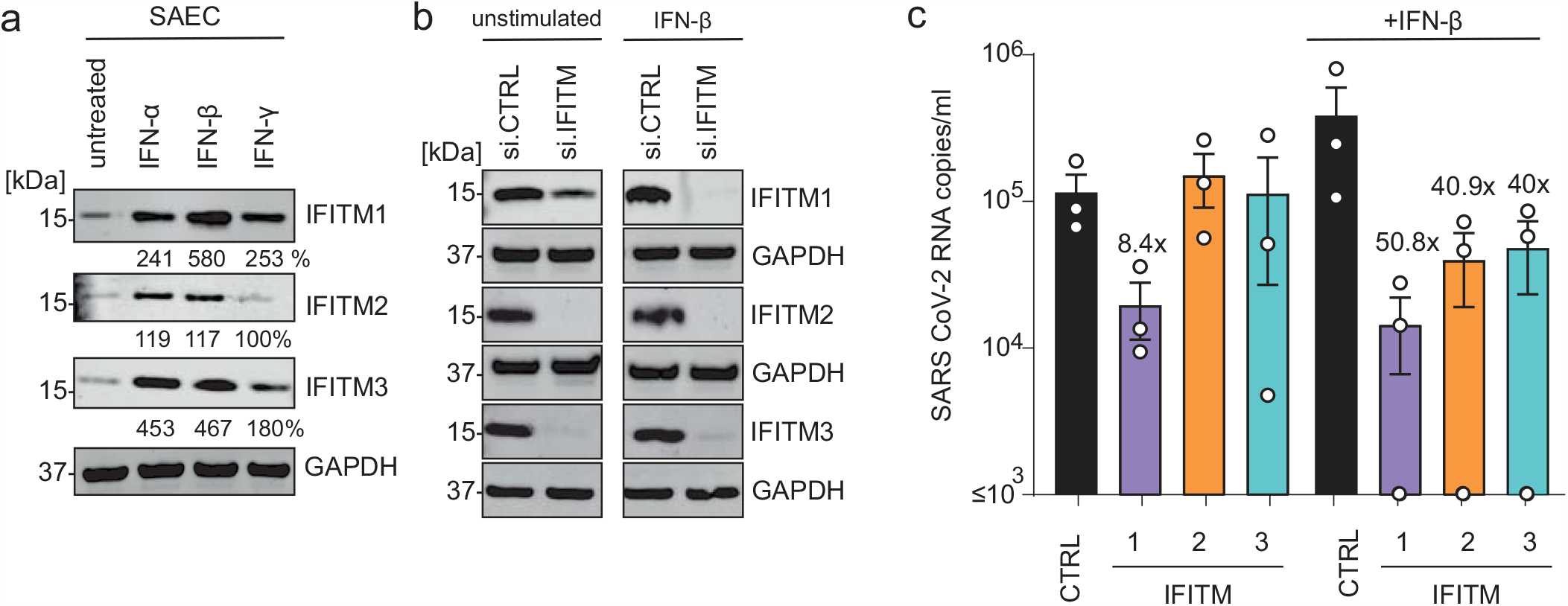
Role of IFITMs in SARS-CoV-2 replication in SAEC. **a**, Expression of IFITM1, IFITM2 and IFITM3 in SAEC after stimulation with IFN-α2 (500 U/ml, 72 h), IFN-β (500 U/ml, 72 h) or IFN-γ (200 U/ml, 72 h). Immunoblots of whole cell lysates were stained with anti-IFITM1, anti-IFITM2, anti-IFITM3 and anti-GAPDH. **b**, Expression of IFITM proteins in SAEC treated with non-targeting or IFITM specific siRNAs. Cells were either stimulated with IFN-β (500 U/ml, 72 h) or left untreated. Immunoblots of whole cell lysates were stained with anti-IFITM1, anti-IFITM2, anti-IFITM3 and anti-GAPDH. **c**, SARS-CoV-2 N quantification in the supernatant of SAEC 2 days post-infection with SARS-CoV-2 (MOI 2.5).

To approximate the *in vivo* situation, we also examined the role of endogenous IFITM expression on genuine SARS-CoV-2 infection of human lung cells. In striking contrast to the results obtained with pseudovirions and/or IFITM overexpression, silencing of endogenous IFITM expression in Calu-3 cells strongly impaired viral RNA production (Fig. 1d, Extended Data Fig. 2c-e). On average, IFITM2 reduced viral RNA yields by ~20-fold in the absence and by~68-fold in the presence of IFN-β. Consequently, the amount of infectious SARS-CoV-2 particles in the cell culture supernatant was reduced by several orders of magnitude upon silencing of IFITM2 and to a lesser extent also by depletion of IFITM1 and IFITM3 (Fig. 1e). Titration analyses showed that IFITMs do not promote SARS-CoV-2 infection in transfected HEK239T cells over a broad range of expression levels (Extended Data Fig. 3). Thus, the opposing effects of transient and endogenous IFITM expression were not just due to different expression levels.

**Fig 3.**
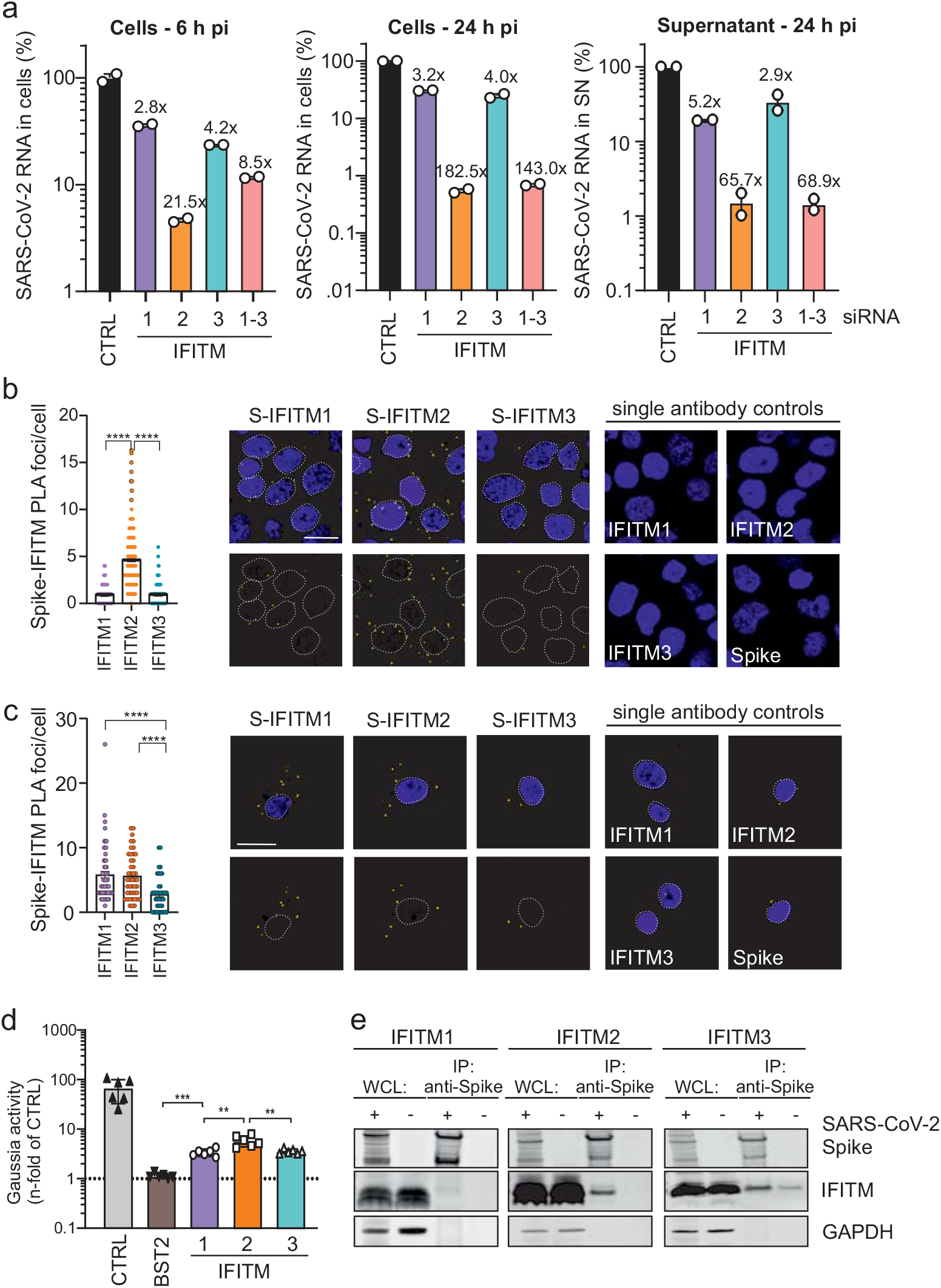
IFITM2 promotes SARS-CoV-2 entry and interacts with the Spike protein. **a**, Intracellular RNA containing N gene sequences copy numbers in Calu-3 cells 6 h (left) and 24 h (middle) post-infection with SARS-CoV-2 (MOI 0.05). Values were normalized to GAPDH and calculated relative to the control (set to 100%). The right panel shows viral RNA copies in the cell culture supernatant at 24 h post infection. Cells were transiently transfected with siRNA either control (CTRL) or targeting IFITM1, 2, 3, or a combination of the three as indicated. Bars represent n=1, measured in duplicates, ±SD. **b**, Proximity ligation assay between the SARS-CoV-2 Spike and IFITM proteins in Calu-3 cells infected with SARS-CoV-2 for 2 h at 4°C. DAPI (blue), nuclei. PLA signal (yellow), proximity between S/IFITMs. Results represent two independent experiments done in technical duplicates. **c**, PLA in SAEC. Bars represent means of n=1 (45-70 cells) ±SEM. DAPI (blue), nuclei. PLA signal (yellow), proximity between S/IFITMs. Scale bar, 20 µm. **d**, Relative interaction between SARS-CoV-2 Spike and human IFITM proteins measured by MaMTH protein-protein interaction assay in cotransfected HEK293T B0166 *Gaussia* luciferase reporter cells. Bars represent the mean of triplicate transfections performed in two independent experiments. **e**, Immunoprecipitation of IFITM proteins by the Spike protein. HEK293T cells were transfected with or without a construct to overexpress SARS-CoV-2 S (indicated with a + or a −) and IFITM1, IFITM2 or IFITM3. 24 h post transfection, cells were harvested and SARS-CoV-2 Spike was immunoprecipitated. WCL, whole cell lysates.

### IFITMs enhance SARS-CoV-2 infection of primary human lung cells

To confirm that the requirement of endogenous IFITM expression for efficient SARS-CoV-2 replication is not limited to Calu-3 cells, we silenced IFITM proteins in primary small airway epithelial cells (SAEC) isolated from normal human lung tissues. Western blot analyses showed that SAEC cells express all three IFITM proteins and type I or II IFN treatment enhanced the expression levels ~2-5-fold (Fig. 2a). siRNA-mediated silencing strongly reduced the expression of IFITM proteins (Fig. 2b) and was associated with ~40-to 50-fold lower levels of SARS-CoV-2 RNA production in the presence of IFN-β (Fig. 2c). Silencing of IFITM1 also clearly reduced viral RNA yields in the absence of IFN treatment (Fig. 2c). Altogether, IFITM1 was more critical for efficient SARS-CoV-2 replication in SAEC cells than in Calu-3 cells (Figs. 1d, 2c). It is thought that IFITM1 is mainly found at the cell surface, while IFITM2 is preferentially localized in early endosomes^6,7^. SARS-CoV-2 may enter cells at their surface as well as in endosomes^17^. Thus, together with differences in the expression levels of specific IFITM proteins, cell-type-dependent differences in the major sites of viral fusion may explain differences in the relative dependency of SARS-CoV-2 on endogenous IFITM1 or IFITM2 expression. In contrast to the results obtained in Calu-3 cells (Extended Data Fig. 2e), IFN-β enhanced rather than inhibited SARS-CoV-2 replication in SAEC cells (Fig. 2c). While this finding came as surprise, it is reminiscent of previous data showing that IFN treatment promotes infection by human coronavirus HCoV-OC43. Notably, this CoV was proposed to hijack IFITM3 for efficient entry^18^. Taken together, our results show that endogenous expression of IFITM proteins promotes SARS-CoV-2 replication in primary human lung cells, especially in the presence of IFN.

### Endogenous IFITMs promote an early step of SARS-CoV-2 infection

To address the mechanisms underlying these opposing effects of IFITMs, we examined the effect of IFITM proteins on SARS-CoV-2 S-mediated fusion under various conditions. To analyse the impact of IFITMs on S-mediated fusion between virions and target cells, we used HIV-1 particles containing β-lactamase-Vpr fusions as previously described^19^, except that the virions contained the SARS-CoV-2 S instead of the HIV-1 Env protein. In agreement with the documented role of IFITMs as inhibitors of viral fusion^12,14^, transient overexpression of all three IFITM proteins blocked fusion of SARS-CoV-2 S HIVpp^19^ with ACE2 expressing HEK293T cells (Extended Data Fig. 4a). Consistent with recent data^20^, results from a split-GFP assay showed that artificial overexpression of IFITMs also inhibits HEK293T cell-to-cell fusion mediated by the SARS-CoV-2 S protein and the ACE2 receptor (Extended Data Fig. 4b). To analyse the impact of endogenous IFITM expression on genuine SARS-CoV-2 entry, we determined the levels of viral RNA in the cells at different time points after infection of Calu-3 cells. Already at 6 h post-infection, depletion of IFITMs 1, 2 and 3 reduced the levels of viral RNA in the cells about 3-, 22- and 4-fold, respectively (Fig. 3a). At 24 h post-infection, silencing of IFITM2 expression decreased intracellular SARS-CoV-2 RNA levels by 182.5-fold and extracellular viral RNA yield by 65.7-fold (Fig. 3a). These results support that in striking contrast to the overexpressed proteins, endogenous IFITM expression is required for efficient SARS-CoV-2 entry into human lung cells.

**Fig 4.**
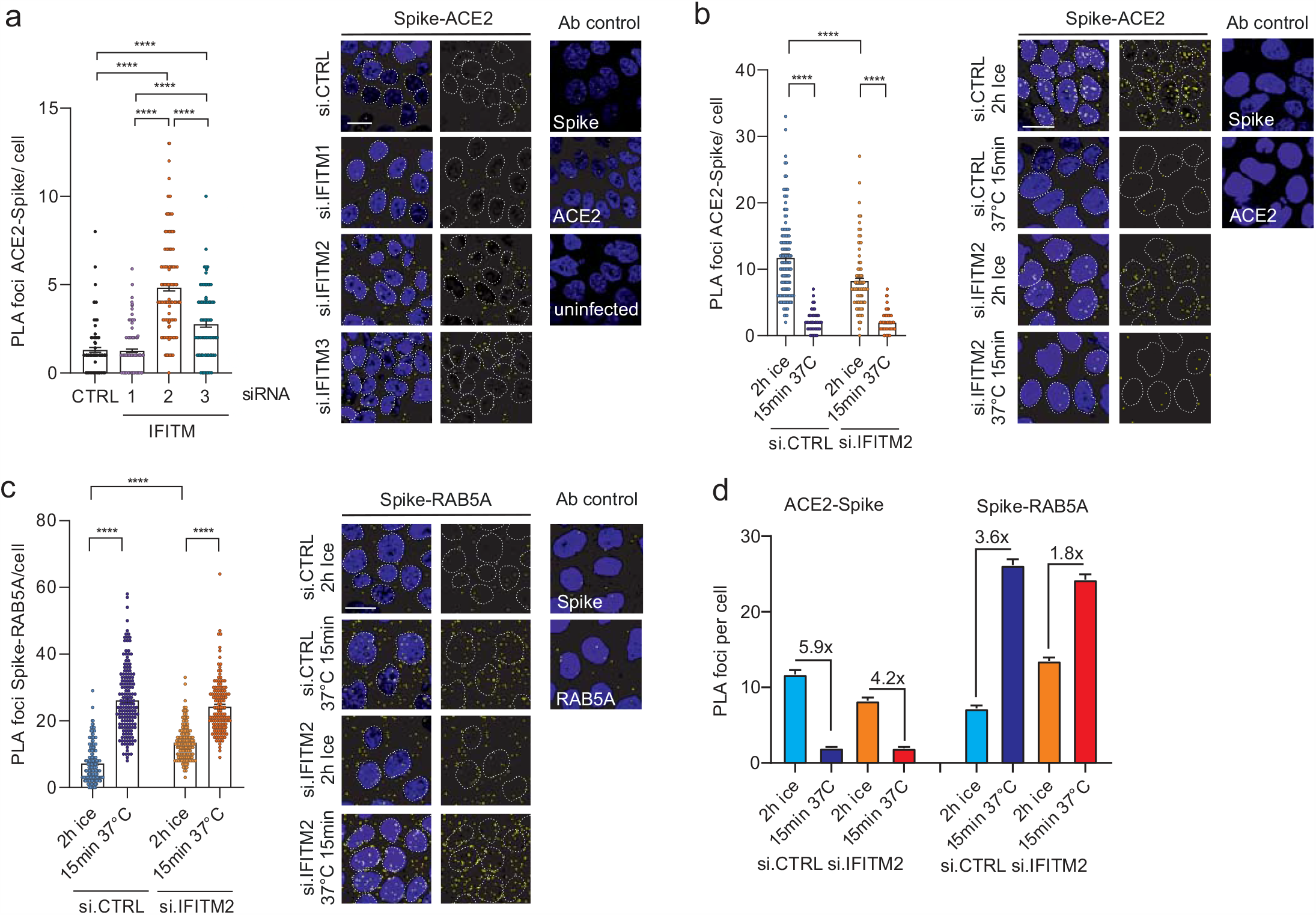
Impact of IFITMs on the ACE2-SARS-CoV-2 S proximity. **a**, PLA between SARS-CoV-2 Spike and ACE2 in Calu-3 depleted of IFITM1, IFITM2 or IFITM3 and infected with genuine SARS-CoV-2. Lines represent means of n=2 (a) n=3 (b) (60-100 cells) ±SEM. **b**, PLA between Spike and ACE2 in Calu-3 cells depleted of IFITM2 and infected with SARS-CoV-2 virus on ice for 2 h and then incubated for 15 min at 37°C. Lines represent means of n=3 (200-300 cells) ±SEM. **c**, PLA assay between Spike and RAB5A in Calu-3 cells infected as in **c**. Lines represent means of n=2 (130-200 cells) ±SEM. DAPI (blue), nuclei. PLA signal (yellow). Scale bar, 20 µm. **d**, Quantification of ACE2-Spike and Spike-RAB5 alpha proximity upon SARS-CoV-2 infection.

### The SARS-CoV-2 Spike interacts with IFITM proteins

It is thought that the broad-spectrum antiviral activity of IFITM proteins does not involve specific interactions with viral proteins but effects on the properties of cellular membranes^5,7,21^. To assess whether the ability of SARS-CoV-2 to utilize IFITMs for efficient infection of human lung cells may instead involve specific interactions between the viral S protein and IFITMs, we performed proximity ligation assays (PLA; Extended Data Fig. 5)^22^. The result revealed higher number of foci for S and IFITM2 compared to IFITM1 and 3 in SARS-CoV-2 infected Calu-3 cells (Fig. 3b), indicating close proximity of these two proteins. In accordance with the relevance of IFITM1 for SARS-CoV-2 replication in this cell type (Fig. 2c), high levels of PLA signals were detected for S and IFITM1 in infected SAEC cells (Fig. 3b). Assessing integral membrane protein-protein interactions using the mammalian-membrane two-hybrid (MaMTH) assay^23^ provided further evidence that SARS-CoV-2 S interacts with IFITM proteins (Fig. 2d, Extended Data Fig. 6). Finally, the SARS-CoV-2 S-protein co-immunoprecipitated IFITM2 and, to a lesser extent, IFITM1 and IFITM3 (Fig. 2e). Altogether, several independent lines of evidence support that the S protein of SARS-CoV-2 interacts with human IFITM proteins.

**Fig 5.**
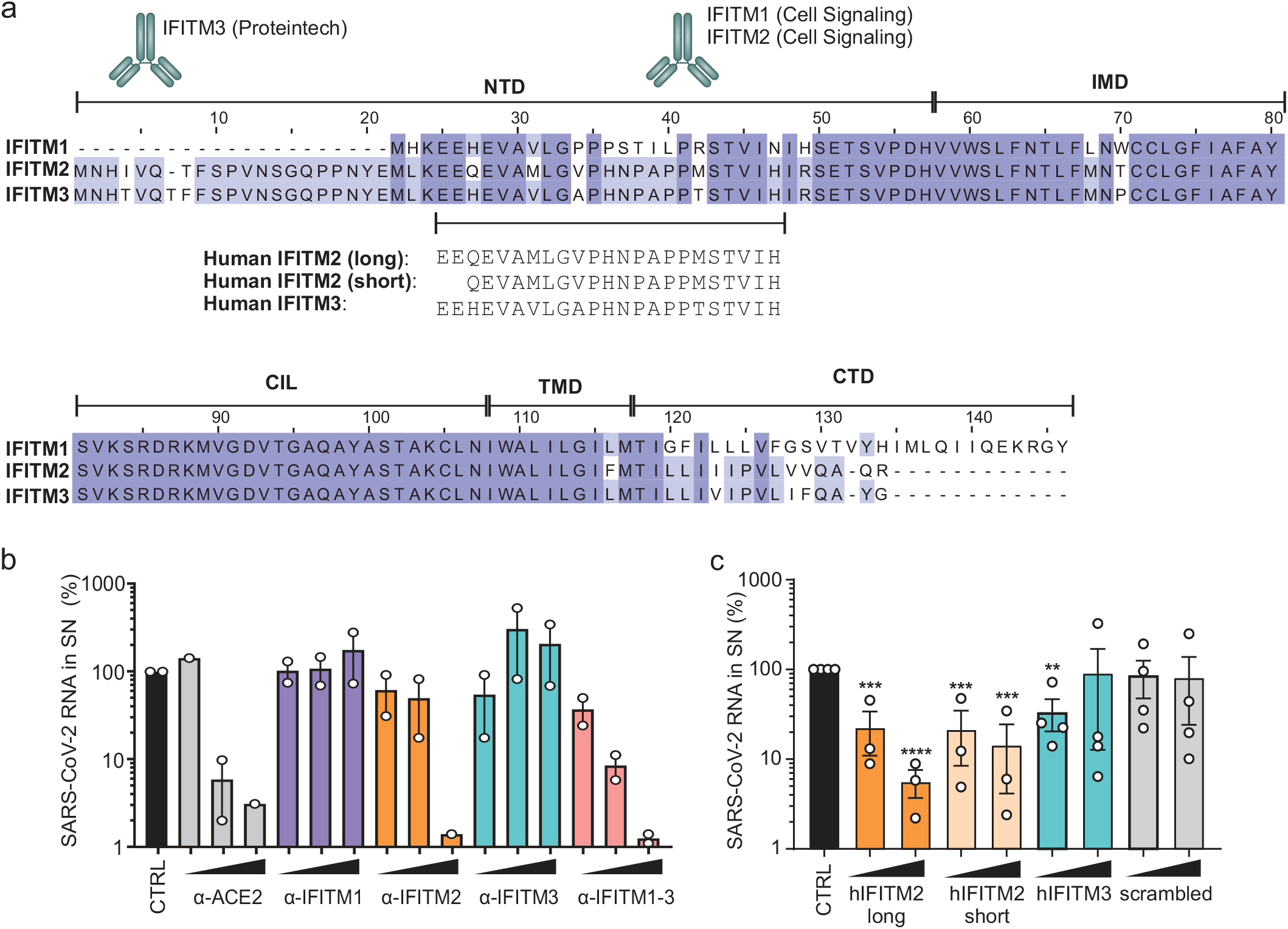
IFITM blocking antibodies and IFITM derived peptides target the N-terminal domain. **a**, Alignment of the amino acid sequence of human IFITM1, 2 and 3. Binding sites of IFITM blocking antibodies are indicated and the region of origin of the IFITM derived peptides highlighted. **b**, Viral N gene RNA levels in the supernatant of Calu-3 cells treated with α-ACE2, α-IFITM1, α-IFITM2, α-IFITM3 and α-IFITM1-3 antibodies, collected 48 h post infection (MOI 0.05). Bars represent one to two independent experiments each measured in technical duplicates (mean value, ±SEM). **c**, RNA containing N gene sequences in the supernatant of Calu-3 cells treated with IFITM-derived peptides, collected 48 h post infection (MOI 0.05). Bars represent two to three independent experiments each measured in technical duplicates (mean value, ±SEM).

**Fig 6.**
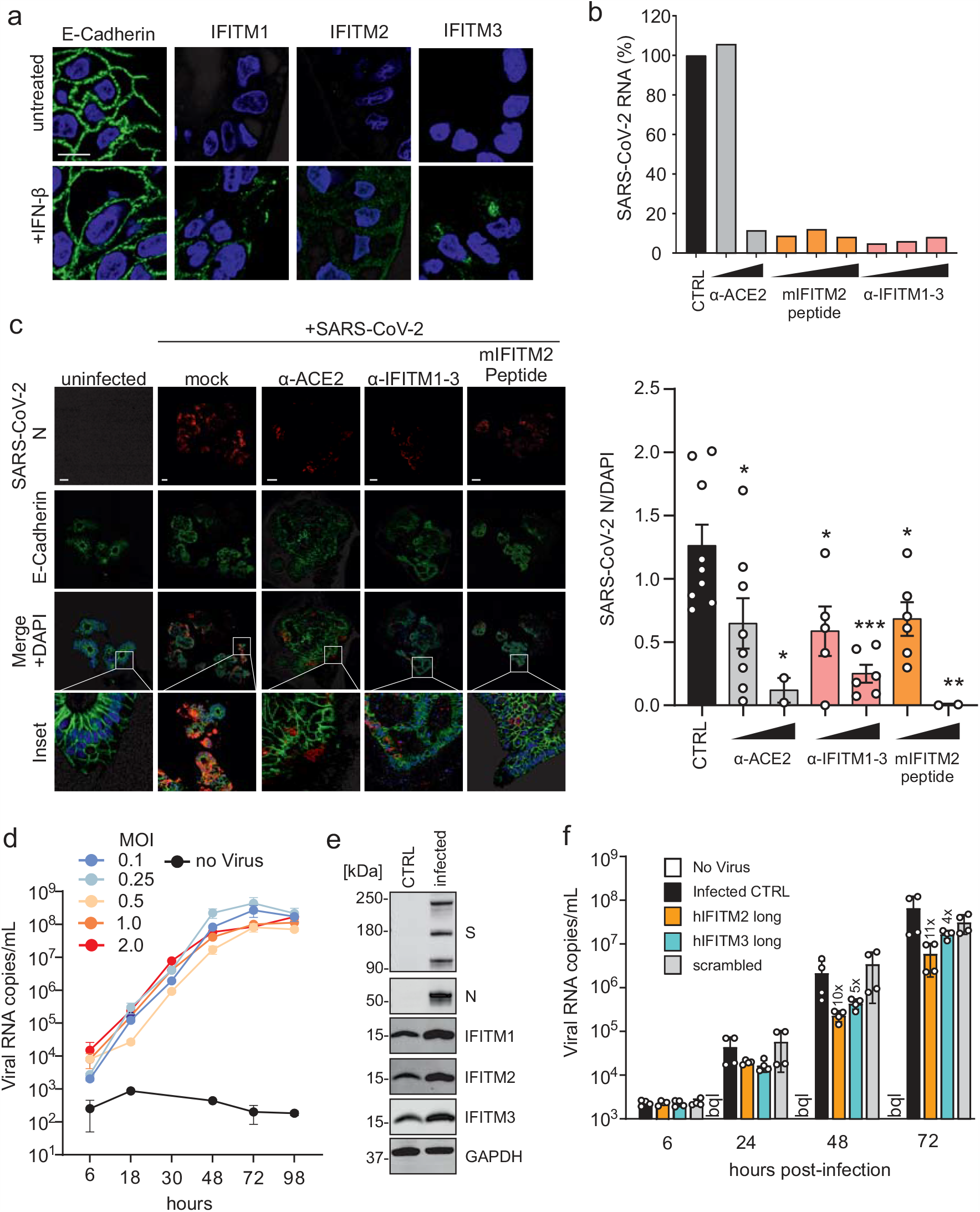
Blocking antibodies and IFITM-derived peptides treatment decrease SARS-CoV-2 infection in gut organoids and cardiomyocytes. **a**, Immunofluorescence images of stem cell-derived gut organoids after stimulation with IFN-β (500 U/ml, 72 h) **b**, Cell-associated viral N gene RNA copy numbers in organoids treated with α-ACE2, mIFITM2 antibody blocking peptide and α-IFITM1-3 and infected with SARS-CoV-2 (MOI 0.15).**c**, Immunohistochemistry of gut organoids treated as in **e** and infected with SARS-CoV-2 (MOI 0.5). Organoids were stained with anti SARS-CoV-2 N (red), E-Cadherin (green) and DAPI (blue). Scale bar, 100 µm (left panel). SARS-CoV-2 N quantification of infected gut organoids treated as in **e** (right panel). **d**, Viral N gene RNA levels in the supernatant of SARS-CoV-2 infected cardiomyocytes (increasing MOIs as indicated), virus containing supernatants at indicated timepoints. **e**, Expression of IFITM1, IFITM2 and IFITM3 in cardiomyocytes infected with SARS-CoV-2. Immunoblot of whole cell lysates stained with anti-IFITM1, anti-IFITM2, anti-IFITM3 and anti-GAPDH **f**, Viral N gene RNA levels in the supernatant of SARS-CoV-2 infected cardiomyocytes (0.05 MOI) treated with IFITM-derived peptides, collected at indicated timepoints post infection. Bars represent two independent experiments each measured in technical duplicates (mean value, ±SEM). bql, below quantification level.

### Effects of endogenous IFITM expression on Spike-ACE2 interaction

Next, we examined whether IFITMs affect the interaction between the SARS-CoV-2 S protein and the ACE2 receptor. Knockdown of IFITM2 and, to a lesser extent, IFITM3 enhanced the number of S/ACE2 PLA foci after infection of Calu-3 cells with genuine SARS-CoV-2 (Fig. 4a). The number of S/ACE2 foci rapidly declined (Fig. 4b) and S/RAB5A signals strongly increased (Fig. 4c) after switching SARS-CoV-2 infected Calu-3 cell cultures from ice to 37°C, most likely indicating S-mediated virion fusion in endosomes. The magnitude of these effects was reduced upon silencing of IFITM2 expression (Fig. 4d) and endogenous IFITM expression usually decreased the number of S molecules that are in close proximity to the ACE2 receptor. It is tempting to speculate that IFITMs reduce the number of S/ACE2 signals by accelerating virion fusion and hence the disappearance of signals. However, further studies are required to elucidate the details of the underlying mechanism(s).

### IFITMs are targets for inhibition of SARS-CoV-2 replication

Our discovery that IFITMs serve as cofactors for efficient SARS-CoV-2 infection suggested that they might represent targets for viral inhibition. To address this, we examined the effect of antibodies targeting the N-terminal region of the three IFITM proteins (Fig. 5a) on SARS-CoV-2 infection of Calu-3 cells. Indeed, antibodies against the N-terminal region of IFITM2 or recognizing all three IFITM proteins inhibited SARS-CoV-2 replication in Calu-3 cells up to 50-fold, while antibodies against IFITM1 or IFITM3 had negligible inhibitory effects (Fig. 5b). Since the membrane topology of IFITMs proteins is under debate^7^, we verified by flow cytometry analyses that the N-terminal region of IFITMs is accessible to antibody binding (Extended Data Fig. 7). Further analyses showed that peptides corresponding to the N-proximal region of IFITM2 that is recognized by inhibitory antibodies also efficiently impair SARS-CoV-2 replication (Fig. 5c). In contrast, the corresponding IFITM3-derived peptide, which differs in four of the 23 residues from the IFITM2-derived peptide, and a scrambled control peptide of the same length and amino acid composition had little if any effect on viral RNA yields. Notably, incubation of SARS-CoV-2 virions with the peptides prior to infection had no inhibitory effect (Extended Data Fig. 8). Thus, similarly to other inhibitors of SARS-CoV-2 infection^24,25^ the IFITM2-derived peptides might target a region in the viral S protein that only becomes accessible during the entry process.

### IFITM-derived peptides or targeting antibodies protect gut organoids and cardiomyocytes against SARS-CoV-2

To better assess the potential relevance of IFITMs for viral spread and pathogenesis in SARS-CoV-2-infected individuals, we analysed their expression in various cell types. We found that IFITM proteins are efficiently expressed in primary human lung bronchial epithelial (NHBE) cells, neuronal cells, and intestinal organoids derived from pluripotent stem cells (Extended Data Fig. 9a-c). These cell types and organoids represent the sites of SARS-CoV-2 entry and subsequent spread, i.e. the lung and the gastrointestinal tract^26–28^, and the potential targets responsible for neurological manifestations of COVID-19^29^. Confocal microcopy analyses confirmed efficient induction of IFITM expression by IFN-β (Fig. 6a). NHBE cells and cultures of neuronal cells did not support efficient SARS-CoV-2 replication precluding meaningful inhibition analyses. Gut organoids, however, are susceptible to SARS-CoV-2 replication^27^ and treatment with the IFITM2-derived peptide or an antibody targeting the N-terminus of IFITMs strongly reduced viral RNA production (Fig. 6b). Independent infection experiments confirmed that both agents significantly reduce viral N protein expression and cytopathic effects in gut organoids (Fig. 6c). Following up on recent evidence that SARS-CoV-2 causes cardiovascular disease^30^, we investigated viral replication in human iPSC-derived cardiomyocytes. In agreement with published data^31^, beating cardiomyocytes were highly susceptible to viral replication (Fig. 6d). All three IFITM proteins were expressed in cardiomyocytes and further induced by virus infection (Fig. 6e). On average, treatment of cardiomyocytes with the IFITM2- or 3-derived peptides reduced the efficiency of SARS-CoV-2 replication by ~10- and 5-fold, respectively (Fig. 6f). In addition, treatment with these peptides suppressed or prevented disruptive effects of virus infection on the ability of cardiomyocytes to beat in culture. Thus, IFITMs can be targeted to inhibit SARS-CoV-2 replication in cells from various human organs, including the lung, gut and heart.

## Discussion

The present study demonstrates that endogenous expression of IFITMs is required for efficient replication of SARS-CoV-2 in human lung cells. In addition, we show that IFITMs can be targeted to inhibit SARS-CoV-2 infection of human lung, gut and heart cells. These findings came as surprise since IFITMs have been reported to inhibit SARS-CoV, MERS-CoV and, very recently, SARS-CoV-2 S-mediated infection^11,15,32^. Confirming and expanding these previous studies, we show that artificial overexpression of IFITM proteins in HEK293T cells prevents S-mediated VSVpp and HIVpp fusion as well as genuine SARS-CoV-2 entry. However, exactly the opposite was observed for genuine SARS-CoV-2 upon manipulation of endogenous IFITM expression in human lung cells: silencing of all three IFITM proteins reduced SARS-CoV-2 entry. Our results provide novel and highly unexpected insights into the role of IFITM proteins in the spread and pathogenesis of SARS-CoV-2 and suggest that these supposedly antiviral factors are hijacked by SARS-CoV-2 as cofactors for efficient entry.

While wildtype IFITM proteins have generally been described as inhibitors of SARS and MERS coronaviruses (Ref) specific point mutations may convert IFITM3 from an inhibitor to an enhancer Spike-mediated pseudoparticle transduction^33^. It has been reported that overexpression of IFITM3 promotes infection by hCoV-OC43, one of the causative agents of common colds^18^. However, IFITM3 was least relevant for SARS-CoV-2 infection in the present study. Thus, although both human coronaviruses may highjack IFITMs for efficient infection they show distinct preferences for specific IFITM proteins. It is under debate whether SARS-CoV-2 mainly fuses at the cell surface or in endosomes and cell-type-specific differences may explain why IFITM2 plays a key role in Calu-3 cells, while IFITM1 is at least as important in SAEC cells. Most importantly, our results clearly demonstrate that IFITM proteins act as critical cofactors for efficient SARS-CoV-2 infection under the most physiological conditions.

We currently do not yet understand why overexpressed and endogenous IFITM proteins have opposite effects on SARS-CoV-2 infection. However, artificial overexpression may change the topology, localisation and endocytic activity of proteins and it has been reported that specific mutations in IFITM3 affecting these features may convert IFITM3 from an inhibitor to an enhancer of coronavirus infection^9,34^. The antiviral activity of IFITMs is very broad and does not involve interactions with specific viral glycoproteins^6,7^. In contrast, the ability of SARS-CoV-2 to hijack IFITMs for efficient entry seems to involve specific interactions between the N-terminal region of IFITMs and the viral S protein (outlined in Extended Data Fig. 10).

IFITMs are strongly induced during the innate immune response in SARS-CoV-2-infected individuals^35,36^. Thus, utilization of IFITMs as infection cofactors may promote SARS-CoV-2 invasion of the lower respiratory tract as well as spread to secondary organs especially under inflammatory conditions. Further studies are required but efficient expression in neurons and cardiomyocytes suggest that IFITMs may play a role in the well documented neuronal and cardiovascular complications associated with SARS-CoV-2 infection (Ref). Perhaps most intriguingly, we show that IFITM-derived peptides and antibodies against the N-terminal region of IFITM2 efficiently inhibit SARS-CoV-2 replication. Targeting cellular IFITM proteins as a therapeutic approach should reduce the risk of viral resistance and be well tolerated since these factors are mainly known for their antiviral activity and may not exert critical physiological functions.

## Methods

### Cell culture

All cells were cultured at 37°C in a 5% CO_2_ atmosphere. Human embryonic kidney 293T cells (HEK293T; ATCC) were maintained in Dulbecco’s Modified Eagle Medium (DMEM) supplemented with 10% heat-inactivated fetal calf serum (FCS), L-glutamine (2 mM), streptomycin (100 µg/ml) and penicillin (100 U/ml). HEK293T were provided and authenticated by the ATCC. Caco-2 (human epithelial colorectal adenocarcinoma) cells were maintained in DMEM containing 10% FCS, glutamine (2 mM), streptomycin (100 µg/ml) and penicillin (100 U/ml), NEAA supplement (Non-essential amino acids (1 mM)), sodium pyruvate (1 mM). Calu-3 (human epithelial lung adenocarcinoma) cells were cultured in Minimum Essential Medium Eagle (MEM) supplemented with 10% FCS (during viral infection) or 20% (during all other times), penicillin (100 U/ml), streptomycin (100 µg/ml), sodium pyruvate (1 mM), and NEAA supplement (1 mM). Hybridoma cells (Mouse I1 Hybridoma CRL-2700; ATCC) were cultured in Roswell Park Memorial Institute (RPMI) 1640 medium supplemented with 10% FCS, L-glutamine (2 mM), streptomycin (100 µg/ml) and penicillin (100 U/ml). Vero cells (ATCC, CCL-81) cells were maintained in DMEM containing 2.5% FCS, glutamine (2 mM), streptomycin (100 µg/ml) and penicillin (100 U/ml), NEAA supplement (Non-essential amino acids (1 mM)), sodium pyruvate (1 mM). Monoclonal anti-VSV-G containing supernatant was aliquoted and stored at −20°C. NHBE (primary human bronchial/tracheal epithelial, Lonza) cells were grown in Bronchial Epithelial Cell Growth Basal Medium (BEGM, Lonza) and Bronchial Epithelial Cell Growth Medium SingleQuots Supplements and Growth Factors (Lonza). SAEC (Small Airway Epithelial cells, Lonza) were grown in Small Airway Epithelial Cell Growth Basal Medium (SABM, Lonza) and Small Airway Epithelial Cell Growth Medium SingleQuots Supplements and Growth Factors (Lonza).

### Human hESC cultivation and gut organoids differentiation

Human embryonic stem cell (hESC) line HUES8 (Harvard University) was used with permission from the Robert Koch Institute according to the “Approval according to the stem cell law” AZ 3.04.02/0084. Cells were cultured on hESC Matrigel (Corning) in mTeSR1 medium (Stemcell Technologies) at 5% CO_2_ and 37°C. Medium was changed every day and cells were splitted twice a week with TrypLE Express (Invitrogen). Experiments involving human stem cells were approved by the Robert-Koch-Institute (Approval according to the stem cell law 29.04.2020).

### Cardiomyocyte differentiation

Human episomal hiPSCs (#A18945, Thermo Fisher Scientific) at passage 2 were split using TrypLE (#12604-013, Thermo Fisher Scientific) to generate a single cell suspension. 18000 iPS cells were seeded on Geltrex (#A1413302, Thermo Fisher Scientific) matrix coated 12 well plates. 3 days post splitting differentiation protocol into iPS cardiomyocytes using the PSC cardiomyocytes Differentiation Kit (#A29212-01, Thermo) was initiated. Contracting iPSC-derived cardiomyocytes were present 14 days post differentiation initiation.

### Neuronal differentiation

Human iPSC, either generated from keratinocytes as previously described^37^ or commercially purchased from the iPSC Core facility of Cedars Sinai (Los Angeles, California), were cultured at 37°C (5% CO_2_, 5% O_2_) on Matrigel-coated (Corning, 354277) 6-well plates using mTeSR1 medium (Stem Cell Technologies, 83850). Neuronal differentiation was chemically induced by culturing hiPSC colonies in suspension in ultra-low attachment T75 flasks (Corning, 3815), to allow the formation of embryoid bodies (EBs). During the first 3 days of differentiation, cells were cultivated in DMEM/F12 (Gibco, 31331-028) containing 20% knockout serum replacement (Gibco, 10828028), 1% NEAA, 1% β-mercaptoethanol, 1% antibiotic-antimycotic, SB-431542 10 µM (Stemcell Technologies, 72232), Dorsomorphin 1 µM (Tocris, 3093), CHIR 99021 3 µM (Stemcell Technologies, 72054), Pumorphamine 1 µM (Miltenyi Biotec, 130-104-465), Ascorbic Acid 200ng/µL, cAMP 500 µM (Sigma-Aldrich, D0260), 1% supplement (Stemcell Technologies, 05731), 0.5% N2 supplement (Gibco, 17502-284). From the fourth day on, medium was switched to DMEM/F12 added with 24 nM sodium selenite (Sigma-Aldrich, S5261), 16 nM progesterone (Sigma-Aldrich, P8783), 0.08 mg/mL apotransferrin (Sigma-Aldrich, T2036), 0.02 mg/mL, Insulin (Sigma-Aldrich, 91077C), 7.72 μg/mL putrescine (Sigma-Aldrich, P7505), 1%NEAA, 1% antibiotic-antimycotic, 50mg/mL heparin (Sigma-Aldrich, H4783), 10 μg/mL of the neurotrophic factors BDNF (Peprotech, 450-02), GDNF (Peprotech, 450-10), and IGF1 (Peprotech, 100-11), 10 μM SB-431542, 1 μM dorsomorphin, 3 μM CHIR 99021, 1 μM pumorphamine, 150 μM. vitamin C, 1 μM retinoic acid, 500 μM cAMP, 1% Neurocult supplement, 0.5% N2 supplement. After 5 further days, neurons were dissociated to single cell suspension and plated onto μDishes, or 6-well plates (Corning, 353046) pre-coated with Growth Factor Reduced Matrigel (Corning, 356231).

### Expression constructs

Expression plasmids encoding for IFITM1, IFITM2 and IFITM3 (pCG_IFITM1, pCG_IFITM2, pCG_IFITM3 and pCG_IFITM1-IRES_eGFP, pCG_IFITM2-IRES_eGFP and pCG_IFITM3-IRES_BFP) were PCR amplified and subcloned in pCG based backbones using flanking restriction sites XbaI and MluI. pCG_SARS-CoV-2-Spike-IRES_eGFP (humanized), encoding the spike protein of SARS-CoV-2 isolate Wuhan-Hu-1, NCBI reference Sequence YP_009724390.1 while pCG_SARS-CoV-2-Spike C-V5-IRES_eGFP was PCR amplified and subcloned using XbaI+MluI, while pCG_SARS-CoV2-Spike C-V5-IRES_eGFP was PCR amplified and subcloned using XbaI+MluI. To generate the pLV-EF1a-human ACE2-IRES-puro, pTargeT-hACE2 was provided by Sota Fukushi and Masayuki Saijo (National Institute of Infectious Diseases, Tokyo, Japan). The ORF of ACE2 was extracted with MluI and SmaI and then inserted into the MluI-HpaI site of pLV-EF1a-IRES-Puro.

### Pseudoparticle stock production

To produce pseudotyped VSV(luc/GFP)ΔG particles, HEK293T cells were transfected with pCG_SARS-CoV-2-Spike C-V5-IRES_GFP, as previously described^38^. 24 hours post transfection, the cells were infected with VSVΔG(GFP/luc)*VSV-G at an MOI of 1. The inoculum was removed after 1 h. Pseudotyped particles were harvested at 16 h post infection. Cell debris was removed by centrifugation at 2000 rpm for 5 min. Residual input particles carrying VSV-G were blocked by adding 10 % (v/v) of I1 Hybridoma supernatant (I1, mouse hybridoma supernatant from CRL-2700; ATCC) to the cell culture supernatant. To produce pseudotyped HIV-1(fLuc)Δ*env* particles, HEK293T cells were transfected with pCMVdR8.91 (Addgene) and pSEW-luc2 (Promega, # 9PIE665) or pCMV4-BlaM-vpr (Addgene, #21950) as well as pCG_SARS-CoV-2-Spike C-V5-IRES_eGFP using TransIT-LT1 according to the manufacturer’s protocol. Six hours post transfection, the medium was replaced with DMEM containing only 2.5% FCS. The particles were harvested 48 hours post transfection. Cell debris was pelleted by centrifugation at 2000 rpm for 5 min.

### Target cell assay

HEK293T cells were transiently transfected using PEI^38^ with pLV-EF1a-human ACE2-IRES-puro and pCG-IFITM1-IRES_eGFP or pCG-IFITM2-IRES_eGFP or pCG-IFITM3-IRES_BFP. 24 h post transfection, cells were transduced/infected with HIV-1Δ*env*(fLuc)* SARS-CoV-2 S or VSV(luc)ΔG*SARS-CoV-2 S particles. 16 h post infection Luciferase activity was quantified.

### Luciferase assay

To determine viral gene expression, the cells were lysed in 300µl of Luciferase Lysis buffer (Luciferase Cell Culture Lysis, Promega) and firefly luciferase activity was determined using the Luciferase Assay Kit (Luciferase Cell Culture, Promega) according to the manufacturer’s instructions on an Orion microplate luminometer (Berthold).

### Vpr-BlaM fusion assay

HEK293T cells were seeded and transiently transfected using PEI^38^ with pLV-EF1a-human_ACE2-IRES-puro and pCG_IFITM1, pCG_IFITM2 or pCG_IFITM3. 24 hours post transfection, cells were transferred to a 96-well plate. On the next day, cells were infected with 50 µl HIV-1 Δenv (BlaM-Vpr)-*SARS-CoV-2-S particles for 2.5 h at 37 °C, followed by washing with PBS. Cells were detached and stained with CCF2/AM (1 mM) as previously described^39^. Finally, cells were washed and fixed with 4% PFA. The change in emission fluorescence of CCF2 after cleavage by the BlaM-Vpr chimera was monitored by flow cytometry using a FACSCanto II (BD).

### SARS-CoV-2 virus stock production

BetaCoV/Netherlands/01/NL/2020 or BetaCoV/ France/IDF0372/2020 was propagated on Vero E6 infected at an MOI of 0.003 in serum-free medium containing 1 μg/ml trypsin as previously described^16^. Briefly, the cells were inoculated for 2 h at 37°C before the inoculum was removed. The supernatant was harvested 48 h post infection upon visible cytopathic effect (CPE). To remove the debris, the supernatants were centrifuged for 5 min at 1,000 × g, then aliquoted and stored at −80°C. Infectious virus titre was determined as plaque forming units (PFU).

### Plaque-forming Unit Assay

The plaque-forming unit (PFU) assay was performed as previously described^16^. SARS-CoV-2 stocks were serially diluted and confluent monolayers of Vero E6 cells infected. After incubation for 2 h at 37°C with shaking every 20 min. The cells were overlaid with 1.5 ml of 0.8 % Avicel RC-581 (FMC) in medium and incubated for 3 days. Cells were fixed with 4 % PFA at room temperature for 45 min. After the cells were washed with PBS once 0.5 ml of staining solution (0.5 % crystal violet and 0.1 % triton in water). After 20 min incubation at room temperature, the staining solution was removed using water, virus-induced plaque formation quantified, and PFU per ml calculated.

### qRT-PCR

N (nucleoprotein) RNA levels were determined in supernatants or cells collected from SARS-CoV-2 infected cells 6 h, 24 h or 48 h post-infection. Total RNA was isolated using the Viral RNA Mini Kit (Qiagen) according to the manufacturer’s instructions. qRT-PCR was performed according to the manufacturer’s instructions using TaqMan Fast Virus 1-Step Master Mix (Thermo Fisher) and a OneStepPlus Real-Time PCR System (96-well format, fast mode). Primers were purchased from Biomers and dissolved in RNAse free water. Synthetic SARS-CoV-2-RNA (Twist Bioscience) were used as a quantitative standard to obtain viral copy numbers. All reactions were run in duplicates. (Forward primer (HKU-NF): 5’-TAA TCA GAC AAG GAA CTG ATT A-3’; Reverse primer (HKU-NR): 5’-CGA AGG TGT GAC TTC CAT G-3’; Probe (HKU-NP): 5’-FAM-GCA AAT TGT GCA ATT TGC GG-TAMRA). GAPDH primer/probe sets (Thermo Fisher) were used for normalization of cellular RNA levels.

### IFITM1, 2 and 3 knock-down

24 h and 96 h after seeding, Calu-3 or SAEC cells were transfected twice with 20 µM of either non-targeting siRNA or IFITM1, IFITM2 or IFITM3 specific siRNA using Lipofectamine RNAiMAX (Thermo Fisher) according to the manufacturer’s instructions. 14 h post transfection, medium was replaced with fresh medium supplemented with 500 U/ml IFN-β in the indicated conditions. 7 h after the second transfection, Calu-3 or SAEC cells were infected with SARS-CoV-2 with an MOI of 0.05 and 2.5 respectively. 6 h later, the inoculum was removed, cells were washed once with PBS and supplemented with fresh media. 48 h post infection, cells and supernatants were harvested for Western blot and qRT-PCR analysis respectively.

### Stimulation with type I interferon

Calu-3, NHBE cells and SAEC were seeded in 12-well plates. For the gut organoids stimulation, HUES88 were seeded in 24-well-plates were coated with growth factor reduced (GFR) Matrigel (Corning) and in mTeSR1 with 10 µM Y-27632 (Stemcell technologies). The next day, differentiation to organoids was started at 80-90% confluency as previously described^26^. Cells or organoids were stimulated with IFN-α2 (500 U/ml, R&D systems 11100-1), IFN-β (500 U/ml, R&D systems 8499-IF-010) or IFN-γ (200 U/ml, R&D systems 285-IF-100). 3 days post-stimulation whole cell lysates were generated.

### Cardiomyocytes infection and kinetics

Human iPSC-derived cardiomyocytes were cultures in 12 wells plates, until they were 3 to 4 weeks old and homogenously beating. Cells were infected with increasing MOIs (0.1, 0.25, 0.5, 1, 2) of the BetaCoV/Netherlands/01/NL/2020 strain. 6 h post infection, cells were washed once with PBS to remove input virus and supplemented with fresh media. Virus-containing supernatant was harvested every day and replaced with fresh media until day 7 (as indicated). N gene RNA copies were determined by qRT-PCR and cells were harvested for Western blot analysis at the latest timepoint.

### Peptides synthesis

The IFITM-derived peptides were synthetized by UPEP, Ulm using F-moc chemistry. Purification to homogeneity of more than 95% was done by reverse phase HPLC. Peptide stock were prepared in distilled water to a final concentration of 10 mg/ml.

### Inibition by IFITM antibodies and peptides

Calu-3 cells were seeded in 48-well format (peptides assays), or in 24-well format (antibodies assay), 24h later cells were treated with increasing concentrations (20 and 80µg/ml) of IFITMs derived peptides (human IFITM2 long: EEQEVAMLGVPHNPAPPMSTVIH, human IFITM2 short: QEVAMLGVPHNAPPMST-VIH, mouse IFITM2 long: EEYGVTELGEPSNSAVVRTTVIN, human IFITM3 long: EEHEVAVLGAPHNPAPPTSTVIH, scrambled IFITM2: EGESGVTTATVEVVIERNNLPY) or blocking antibodies (15 and 30 µg/ml) (α-ACE2 AK (AC18Z), Santa Cruz Biotechnology sc-73668; α-IFITM1 Cell Signaling 13126 S, α-IFITM2 Cell Signaling 13530S, α-IFITM3 Proteintech 11714-1-AP, α-IFITM1/2/3 (F-12) Santa Cruz Biotechnology sc-374026) as indicated. 2 h post-treatment, cells were infected with SARS-CoV-2 with an MOI of 0.05. 6 h post-infection, cells were washed once with PBS and supplemented with fresh MEM medium. 48 h post-infection supernatants were harvested for qRT-PCR analysis. Cardiomyocytes were seeded in 12-well plates, and treated with 100 µg/ml of indicated peptides 1h prior to infection (MOI 0.01). 6 h post infection, cells were washed once with PBS to remove input virus and supplemented with fresh media. Virus-containing supernatant was harvested every day, replaced with fresh media until day 3, and fresh peptides (100 µg/ml) (as indicated). N gene RNA copies were determined by qRT-PCR. Gut organoids were treated with increasing concentrations (15 and 30 µg/ml) of IFITMs derived peptides (mouse IFITM2 antibody blocking peptide Santa Cruz sc-373676 P) and blocking antibodies (α-ACE2 AK (AC18Z), Santa Cruz Biotechnology sc-73668, α-IFITM1/2/3 (F-12) Santa Cruz Biotechnology sc-374026) as indicated. 1h30 post-treatment, organoids were infected with SARS-CoV-2 with an MOI 0.15 as previously described^40^. 48 h post-infection gut organoids were harvested for qRT-PCR analysis.

### Virus treatment

Calu-3 cells were seeded in 48-wells, 24 h later SARS-COV-2 (0.05 MOI) was incubated for 30 min at 37°C with indicated concentrations of IFITM-derived peptides. 50 µl of the inoculum were used to infect the cells. 6h later cells were supplemented with fresh medium. 48 h post-infection supernatants were harvested for qRT-PCR analysis.

### Flow cytometry analysis of IFITMs

HEK293T cells were transfected with pCG_IFITM1, 2 or 3 using PEI as previously described. Calu-3 cells were seeded 24 h before harvest in a 6 well format. 24h post transfection and post seeding, cells were harvested using a scraper and stained with the eBioscience Fixable Viability Dye eFluor 780 (Thermo Fisher) for 15 minutes at room temperature in the dark. Afterwards cells were washed three times with PBS and fixed with 100µl of Reagent A (FIX & PERM Fixation and Permeabilization Kit, Nordic MUbio) for 30 minutes at room temperature, washed three time with PBS and stained with primary antibody (α-IFITM1 Cell Signaling 13126 S, α-IFITM2 Cell Signaling 13530S, α-IFITM3 Proteintech 11714-1-AP, α-IFITM1/2/3 (F-12) Santa Cruz Biotechnology sc-374026,) diluted 1:20 in PBS or in Reagent B (FIX & PERM Fixation and Permeabilization Kit Nordic MUbio) for 1 h at 4°C. Cells were washed three times with PBS and stained with secondary antibody (Goat Anti-Rabbit IgG H&L (PE), ab72465, Donkey Anti-Mouse IgG H&L (PE) ab7003, 1:50) for 1 h at 4°C. After several washing with PBS, cells were resuspended in 100µl of PBS.

### Immunofluorescence of gut organoids

For histological examination, organoids were fixed in 4 % PFA over night at 4°C, washed with PBS, and pre-embedded in 2 % agarose (Sigma) in PBS. After serial dehydration, intestinal organoids were embedded in paraffin, sectioned at 4 µm, deparaffinized, rehydrated and subjected to heat mediated antigen retrieval in tris Buffer (pH 9) or citrate buffer (pH 6). Sections were permeabilized with 0.5 % Triton-X for 30 min at RT and stained over night with primary antibodies (rabbit anti-IFITM1 Cell Signaling 13126 S, 1:500 or rabbit anti-IFITM2 Cell Signaling #13530S, 1:500 or rabbit anti-IFITM3 Cell Signaling #59212S, 1:250 or anti-SARS-CoV-2 N 1:500 or anti-E-Cadherin 1:500) diluted in antibody diluent (Zytomed) in a wet chamber at 4°C. After washing with PBS-Tween 20, slides were incubated with secondary antibodies (Alexa Fluor IgG H+L, Invitrogen, 1:500) and 500 ng/ml DAPI in Antibody Diluent for 90 min in a wet chamber at RT. After washing with PBS-T and water, slides were mounted with Fluoromount-G (Southern Biotech). Negative controls were performed using IgG controls or irrelevant polyclonal serum for polyclonal antibodies, respectively. Cell borders were visualized by E-cadherin staining. Images were acquired using a LSM 710 system.

### GFP Split fusion assay

GFP1-10 and GFP11-expressing HEK293T cells were seeded separately in a 24-well plate. One day post seeding, cells were transiently transfected using the calcium-phosphate precipitation method ^41^. GFP1-10 cells were co-transfected with increasing amounts (0, 8, 16, 32, 64, 125, 250, 500 ng) of pCG_IFTM1, pCG_IFITM2, pCG_IFITM3 and 250 ng of pLV-EF1a-human ACE2-IRES-puro. GFP11 cells were transfected with 250 ng of pCG_SARS-CoV-2-Spike C-V5 codon optimised. 16 h post transfection, GFP1-10 and GFP11 cells were co-cultured in poly-L-lysine-coated 24-well plate. 24 h post co-culturing, cells were fixed with 4 % PFA and cell nuclei were stained using NucRed Live 647 ReadyProbes Reagent (Invitrogen) according to the manufacturer’s instructions. Fluorescence imaging of GFP and NucRed was performed using a Cytation3 imaging reader (BioTek Instruments). 12 images per well were recorded automatically using the NucRed signal for autofocusing. The GFP area was quantified using ImageJ.

### Whole cell lysates

To determine expression of cellular and viral proteins, cells were washed in PBS and subsequently lysed in Western blot lysis buffer (150 mM NaCl, 50 mM HEPES, 5 mM EDTA, 0.1% NP40, 500 μM Na_3_VO_4_, 500 μM NaF, pH 7.5) supplemented with protease inhibitor (1:500, Roche) as previously described ^38^. After 5 min of incubation on ice, samples were centrifuged (4°C, 20 min, 14.000 rpm) to remove cell debris. The supernatant was transferred to a fresh tube, the protein concentration was measured and adjusted using Western blot lysis buffer. Lysates from iPSC-derived neurons were prepared following previously published protocols^42^. Briefly, neurons were harvested in cold PBS (Gibco) and centrifuged at 5000 RPM for 3 minutes. Pellets were then resuspended and incubated at 4°C on an orbital shaker for 2 hours in RIPA buffer. Lysate were then sonicated and protein concentration was determined by Bradford assay.

### SDS-PAGE and Immunoblotting

Western blotting was performed as previously described^38^. In brief, whole cell lysates were mixed with 4x or 6x Protein Sample Loading Buffer (LI-COR, at a final dilution of 1x) supplemented with 10 % β-mercaptoethanol (Sigma Aldrich), heated at 95°C for 5 min, separated on NuPAGE 4±12% Bis-Tris Gels (Invitrogen) for 90 minutes at 100 V and blotted onto Immobilon-FL PVDF membranes (Merck Millipore). The transfer was performed at a constant voltage of 30 V for 30 minutes. After the transfer, the membrane was blocked in 1 % Casein in PBS (Thermo Scientific). Proteins were stained using primary antibodies against IFITM1 (α-IFITM1, Cell Signaling #13126 S, 1:1000,), IFITM2 (α-IFITM2 Cell Signaling #13530S, 1:1000), IFITM3 (α-IFITM3 Cell Signaling #59212S, 1:1000) SARS Spike CoV-2 (SARS-CoV-1/-2 (COVID-19) spike antibody [1A9], GTX-GTX632604, 1:1000), VSV-M (Mouse Monoclonal Anti-VSV-M Absolute antibody, ABAAb01404-21.0, 1:1000), actin (Anti-beta Actin antibody Abcam, ab8227, 1:5000 Abcam,), ACE2 (Rabbit policclonal anti-ACE2 Abcam, ab166755, 1:1000) and Infrared Dye labelled secondary antibodies (LI-COR IRDye). Membranes were scanned using LI-COR and band intensities were quantified using Image Studio (LI-COR).

### Proximity Ligation Assay

The proximity ligation assay (PLA) was performed as previously described^43^. In brief, Calu-3 or SAEC were seeded in a 24-well plate on a cover slip glass. 24 h and 72 h post seeding, the cells were transfected with 20 µM either non-targeting siRNA or IFITM1 or IFITM3 siRNAs using RNAimax according to the manufacturer’s instructions. Prior infection, cells were pre-chilled for 30 minutes at 4°C and then infected with VSV(luc)ΔG*-SARS-CoV-2 S (MOI 2) or BetaCoV/France/IDF0372/2020 (MOI 0.05) for 2 h on ice. Cells have been washed once with cold PBS and fixed with 4% PFA. For staining following antibodies were used: IFITM1 (α-IFITM1 Cell Signaling 13126 S), IFITM2 (α-IFITM2 Abcam 236735), IFITM3 (α-IFITM3 Cell Signaling 59212S), SARS Spike CoV-2 (SARS-CoV / SARS-CoV-2 (COVID-19) spike antibody [1A9], GTX-GTX632604), Rab5 alpha (Rab5 (RAB5A) Goat Polyclonal Antibody Origene AB0009-200) and ACE2 (Rabbit polyclonal anti-ACE2 Abcam, ab166755). All in a concentration 1:100. Images were acquired on a Zeiss LSM 710 and processed using ImageJ (Fiji).

### Co-immunoprecipitation SARS-CoV-2 Spike and IFITMs

HEK293Ts were transfected using PEI with 0.5 μg pCG-SARS CoV2 Spike-V5 and 0.5 μg of pCG IFITM1, IFITM2 or IFITM3. 24 h later, samples were lysed with IP lysis buffer (50 mM, Tris pH8, 150 mM NaCl, 1 % NP40, protease inhibitor) for 10 min on ice. Lysed samples were centrifuged and incubated for 3 h with Pierce Protein A/G Magnetic beads (88802) which were pre-incubated over night with V5 antibody (Cell signaling E9H80; 5 μg of primary antibody per 10 μl of beads per sample).

### MaMTH assay

Human IFITM proteins and SARS-CoV-2 viral proteins were cloned into MaMTH N-term tagged Prey and C-term tagged Bait vectors respectively using Gateway cloning technology (ThermoFisher). Correctness of recombined insertions was confirmed by Sanger sequencing (Eurofins). The Mammalian Membrane Two-Hybrid (MaMTH) Assay has been performed as previously described^23,44^. HEK293T B0166 Gaussia luciferase reporter cells were co-transfected in 96-well plates with 25 ng SARS-CoV-2 protein Bait and 25 ng IFITM or control protein Prey MaMTH vectors in triplicates using PEI transfection reagent. Gal4 (transcription factor) as well as EGFR Bait with SHC1 Prey served as positive controls, whereas SARS-CoV-2 Bait proteins with Pex7 Prey were used as negative controls. The following day, Bait protein expression was induced with 0.1µg/ml doxycycline. Cell-free supernatants were harvested 2 days post-transfection and the released Gaussia reporter was measured 1 s after injecting 20 mM coelenterazine substrate using an Orion microplate luminometer. To determine the level of protein interaction, Gaussia values were normalized to Pex7 Prey negative control for each Bait. To determine Bait and Prey protein expression levels, HEK293T B0166 transfected and treated in the same manner were harvested two days post-transfection and lysed in Co-IP buffer (150 mM NaCl, 50 mM HEPES, 5 mM EDTA, 0.10% NP40, 0.5 mM sodium orthovanadate, 0.5 mM NaF, protease inhibitor cocktail from Roche) and reduced in the presence of β-mercaptoethanol by boiling at 95°C for 10 min. Proteins were separated in 4 to 12% Bis-Tris gradient acrylamide gels (Invitrogen), blotted onto polyvinylidene difluoride (PVDF) membrane, blocked in 5% milk and probed with rabbit anti-V5 (Cell Signaling #13202), mouse anti-FLAG (Sigma #F1804) and rat anti-GAPDH (Biolegend #607902) antibodies, followed by goat anti-mouse, anti-rabbit and anti-rat secondary fluorescent antibodies (LI-COR). Membranes were scanned with LI-COR Odyssey reader.

### Statistics

Statistical analyses were performed using GraphPad PRISM 8 (GraphPad Software). P-values were determined using a two-tailed Student’s t test with Welch’s correction. Unless otherwise stated, data are shown as the mean of at least three independent experiments ± SEM. Significant differences are indicated as: *, p < 0.05; **, p < 0.01; ***, p < 0.001. Statistical parameters are specified in the figure legends.

## Acknowledgments

We thank K. Regensburger, S. Engelhart, M. Meyer, R. Burger, N. Schrott, N. Preising and D. Krnavek for technical assistance and. The ACE2 vector and the SARS-CoV-2 S-HA plasmid were provided by Shinji Makino and Stefan Pöhlmann. We thank K-K. Conzelmann for providing VSVΔG and the Core Functional Peptidomics of Ulm University for peptide synthesis. This study was supported by DFG grants to F.K., J.Mün., D.K., K.M.J.S., D.Sa. (CRC 1279, SPP 1923, KM 5/1-1, SP1600/4-1), C.G. (GO2153/3-1) EU’s Horizon 2020 research and innovation program to J.M. (Fight-nCoV, 101003555), as well as the BMBF to F.K., D.Sa. and K.M.J.S. (Restrict SARS-CoV-2, protACT and IMMUNOMOD). C.P.B., C.C., and R.G. are part of and R.G. is funded by a scholarship from the International Graduate School in Molecular Medicine Ulm (IGradU).

## Author Contributions

C.P.B. and R.N. performed most experiments. M.V. performed interaction assays. J.K., S.H. and A.K. provided gut organoids. C.M.S. generated most expression constructs. D.K. performed MaMTH assays. J. Mül., C.C. and J. Mün provided SARS-CoV-2. F.Z. assisted in experiments with infectious SARS-CoV-2. L.W., T.W. and R.G. provided reagents and protocols. D.Sc. performed FACS for the Vpr-BlaM assay; E.B. and J.W. performed the HEK293T GFP split fusion assay. L.K. helped with the microscopy analysis of organoids. F.D. and S.J. provided cardiomyocytes. A.C. M.S. and T.B. provided neurons. D.S., C.G., S.S. and J. Mün. provided comments and resources. K.M.J.S and F.K. conceived the study, planned experiments and wrote the manuscript. All authors reviewed and approved the manuscript.

## Data Availability

The datasets generated during and/or analyzed during the current study are available from the corresponding authors on request.

