## Supplementary Files for "IFITM proteins promote SARS-CoV-2 infection and are targets for virus inhibition"

[1]Institute of Molecular Virology, Ulm University Medical Centre, 89081 Ulm, Germany. [2]Department of Internal Medicine I, Ulm University Medical Centre, 89081 Ulm, Germany. [3]Department of Infectious Diseases, King's College London, WC2R 2LS London, United Kingdom. [4]Institute of Medical Virology and Epidemiology of Viral Diseases, University Hospital Tübingen, 72076 Tübingen, Germany. [5]Department of Internal Medicine II (Cardiology), Ulm University, 89081 Ulm, Germany. [6]Institute of Virology, Charité - Universitätsmedizin Berlin, 10117 Berlin, Germany. [7]Institute for Anatomy and Cell Biology, Ulm University, 89081 Ulm Germany. [8]Institute of Medical Microbiology and Hygiene, Ulm University Medical Centre, 89081 Ulm, Germany. [9]Institute of Medical Science, The University of Tokyo, 1088639 Tokyo, Japan.

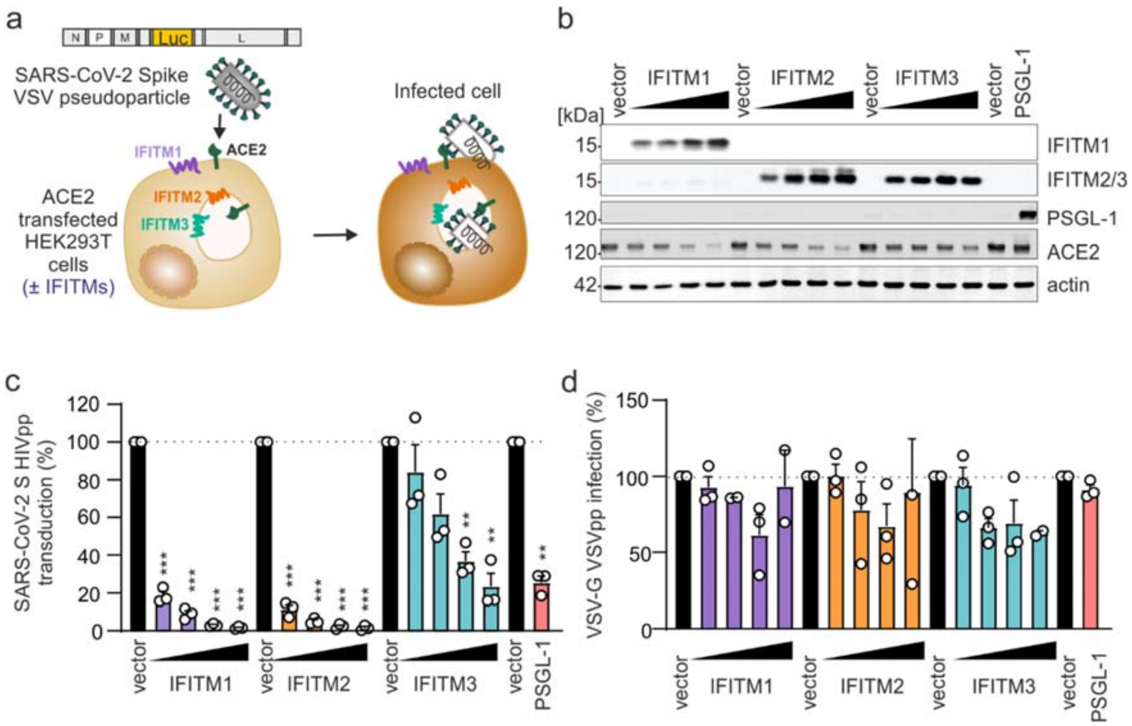


**Extended data Figure 1. Effect of IFITM overexpression on Spike- or VSV-G-mediated pseudoparticle infection. a,** Schematic depiction of the assay to assess VSVpp entry. **b**, Immunoblot of whole cell lysates (WCLs) of HEK293T cells co-transfected with expression plasmids for ACE2, empty or PSGL-1 expressing control vectors, or different doses of IFITM expression constructs. Immunoblots of whole cell lysates were stained with anti-IFITM1, anti-IFITM2, anti-IFITM3 and anti-actin. **c,** Quantification of SARS-CoV-2-S-mediated entry by measuring luciferase activity in HEK293T cells transiently transfected with the indicated expression vectors and transduced 24 h post-transfection with HIV(luc)Δ*env**-SARS-CoV-2 S for 48 h. All bar diagrams in this figure represent means of n=3 (±SEM). PSGL-1 was recently shown to block SARS-CoV-2 attachment[45] and was used as positive control for inhibition. **d,** VSV(luc)ΔG*VSV-G entry in HEK293T cells transiently expressing indicated proteins and infected 24 h post transfection with VSV(luc) ΔG*VSV-G (MOI 0.025) for 16 h. Lower panel: Immunoblot of the corresponding whole cell lysates (WCLs) stained with anti-IFITM1, anti-IFITM2, anti-IFITM3, anti-PSGL-1, anti-ACE2 and anti-actin.

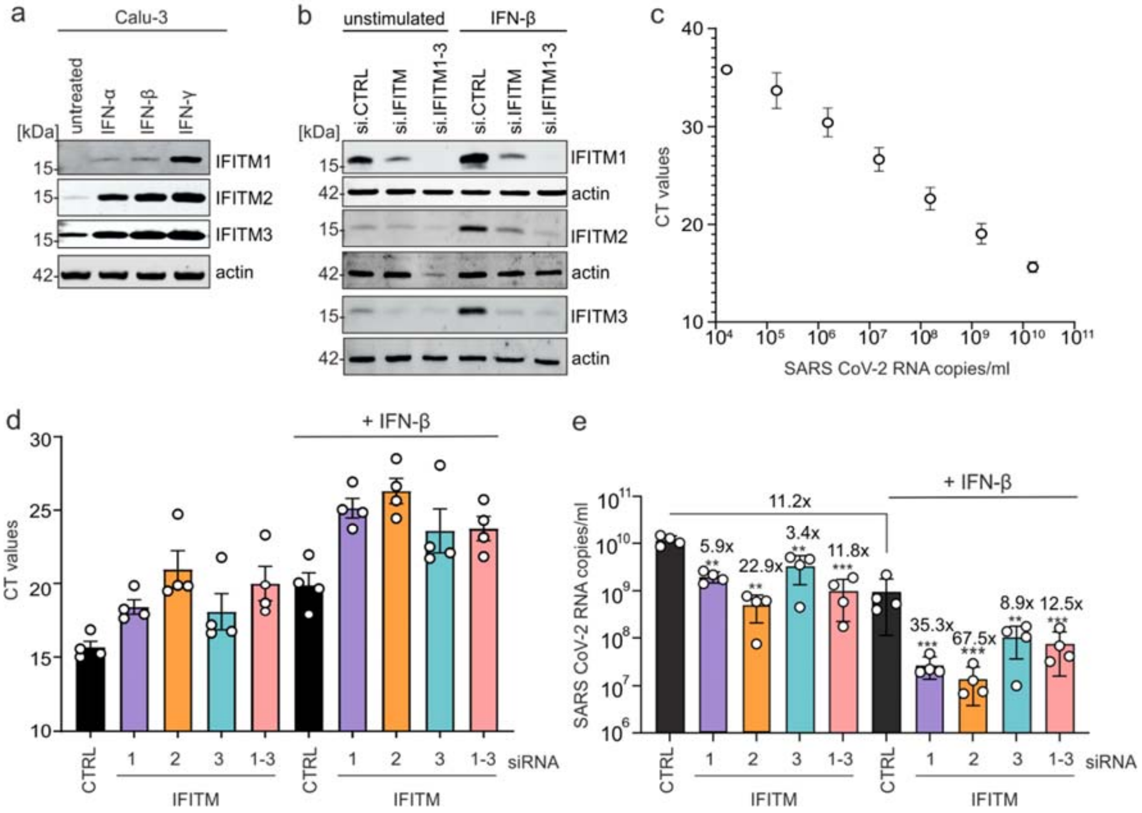


**Extended data Figure 2. Impact of IFITM siRNA knock-down on SARS-CoV-2 replication in Calu-3 cells.** **a**, Expression of IFITM1, IFITM2 and IFITM3 in Calu-3 cells after stimulation with IFN-α2 (500 U/ml, 72 h), IFN-β (500 U/ml, 72 h) or IFN-γ (200 U/ml, 72 h). Immunoblots of whole cell lysates were incubated with anti-IFITM1, anti-IFITM2, anti-IFITM3 and anti-actin. **b**, Expression of IFITM proteins in Calu-3 cells transfected with non-targeting or IFITM-specific siRNAs. Cells were either stimulated with IFN-β (500 U/ml, 72 h) or left untreated. Immunoblots of whole cell lysates were stained with anti-IFITM1, anti-IFITM2, anti-IFITM3 and anti-actin. **c**, Standard curve, **d**, raw qRT-PCR CT values and **e**, SARS-CoV-2 RNA copy numbers in the supernatant of Calu-3 cells collected 2 days post-infection with SARS-CoV-2 (MOI 0.05). Relative levels of viral RNA production and shown in Figure 1d. Number above the bars indicate n-fold reduction of viral RNA levels upon knockdown of the respective IFITM proteins compared to cells treated with control siRNA or fold inhibit by IFN-β treatment, respectively. The bar diagram shows mean values (+/SD) from four independent experiments each measured in technical duplicates.

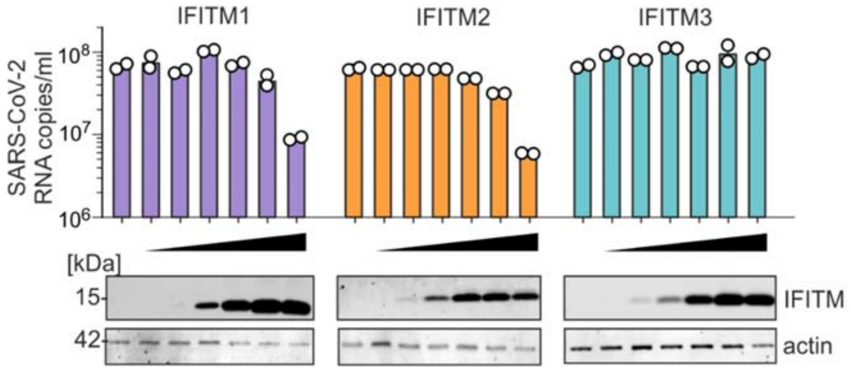


**Extended data Figure 3. Effect of different levels of transient IFITM expression on SARS-CoV-2 infection.** SARS-CoV-2 RNA production from HEK239T cells transiently expressing ACE2 and increasing levels of the indicated IFITM proteins. Quantification of viral N gene RNA by qRT-PCR in the supernatant of HEK293T was performed 48 h post-infection with SARS-CoV-2 (MOI 0.05). Bars represent means of n=2±SEM.

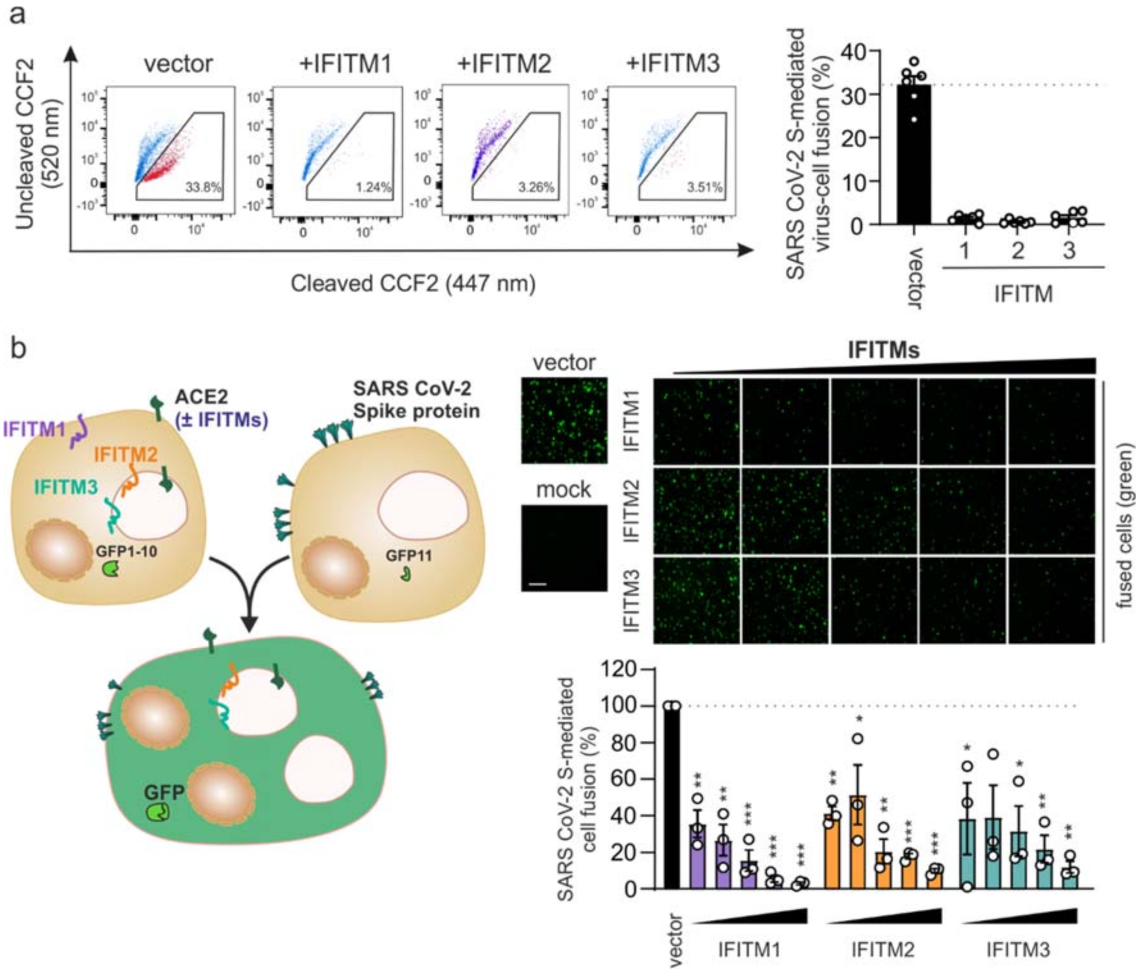


**Extended data Figure 4. Overexpression of IFITMs prevents S-mediated virion and cell-to-cell fusion. a**, Fusion of HIV(Vpr-Blam)Δ*env**-SARS-CoV-2-S with HEK293T cells transiently expressing ACE2 and IFITMs. Quantification of the fusion efficiency by flow cytometry as percentage of (cleaved CCF2) positive cells. Bars represent means (±SEM) of three experiments each done in triplicate. Right panel: Exemplary gating of the raw data. **b**, Schematic outline of the split-GFP assay measuring cell-cell fusion (left). GFP1-11 and SARS-CoV-2 Spike expressing HEK293T were co-cultured with GFP10, ACE2 and IFITM expressing HEK293T. Exemplary fluorescence images (upper). Quantification of successful fusion by GFP positive cells (green) normalized to nuclei (lower right). Bars represent means of n=3, ±SEM.

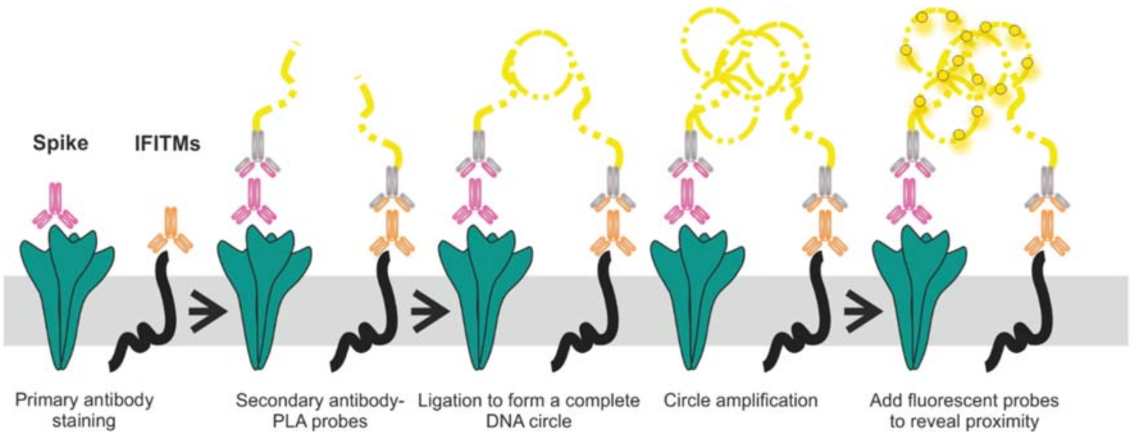


**Extended data Figure 5. Schematic outline of the Proximity ligation assay (PLA).** This assay measures the proximity between SARS-CoV-2 Spike protein and the IFITMs proteins.

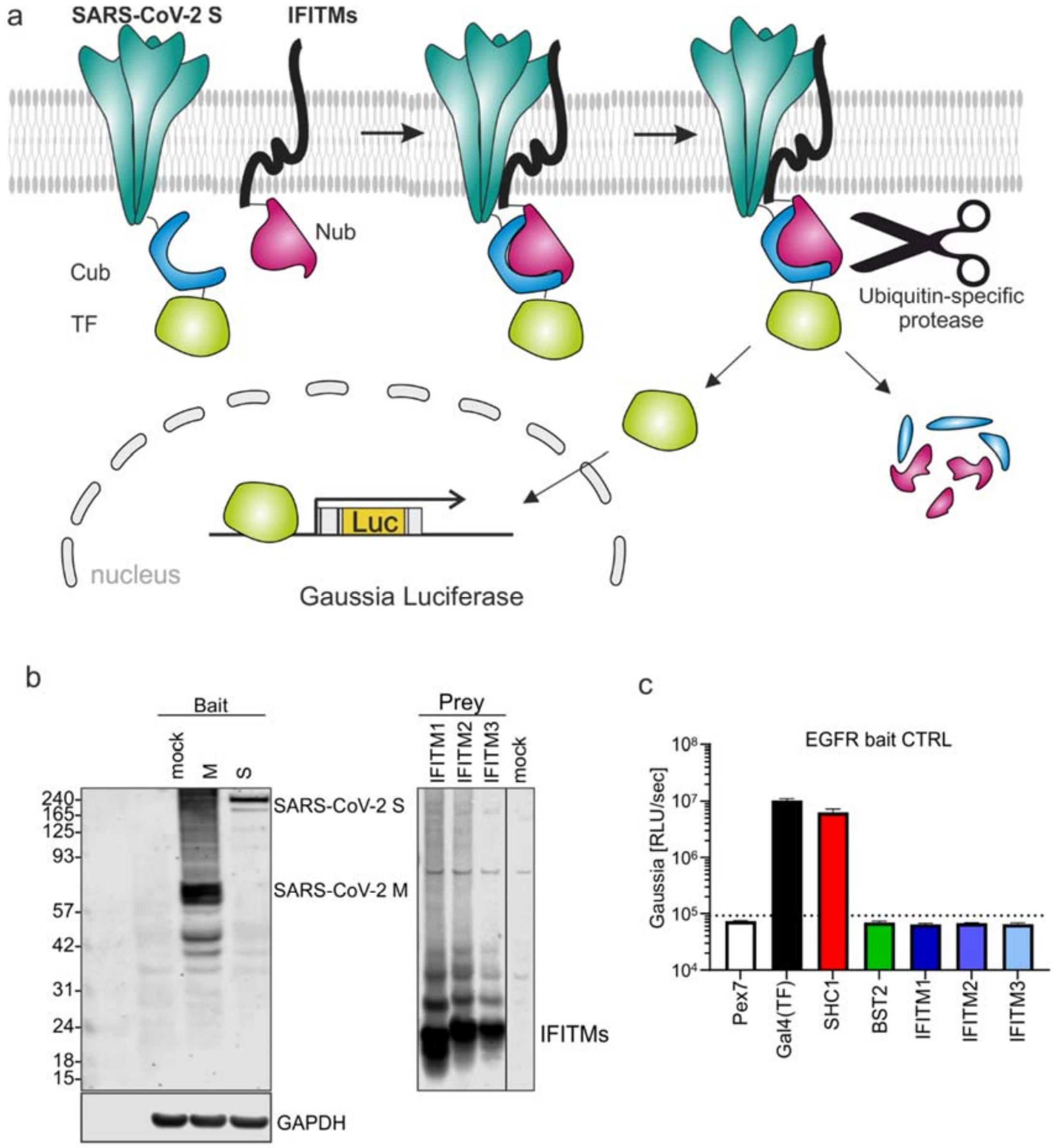


**Extended data Figure 6. Analysis of protein-protein interactions by MaMTH assay. a,** Schematic representation of the MaMTH assay measuring interaction between SARS-CoV-2 Spike and IFITM1, 2 or 3. **b,** Western blot showing the expression of MaMTH V5-tagged SARS-CoV-2 protein Baits and FLAG-tagged IFITM Preys in transfected HEK293T B0166 cells. GAPDH shown as loading control. **c**, Raw values of negative (Baits only or Preys with EGFR Bait) and positive controls (transcription factor Gal4 or EGFR with SHC1) used in MaMTH protein-protein interaction assay. Dotted line indicates untransfected sample value (mock). Mean of triplicate transfection + SD.

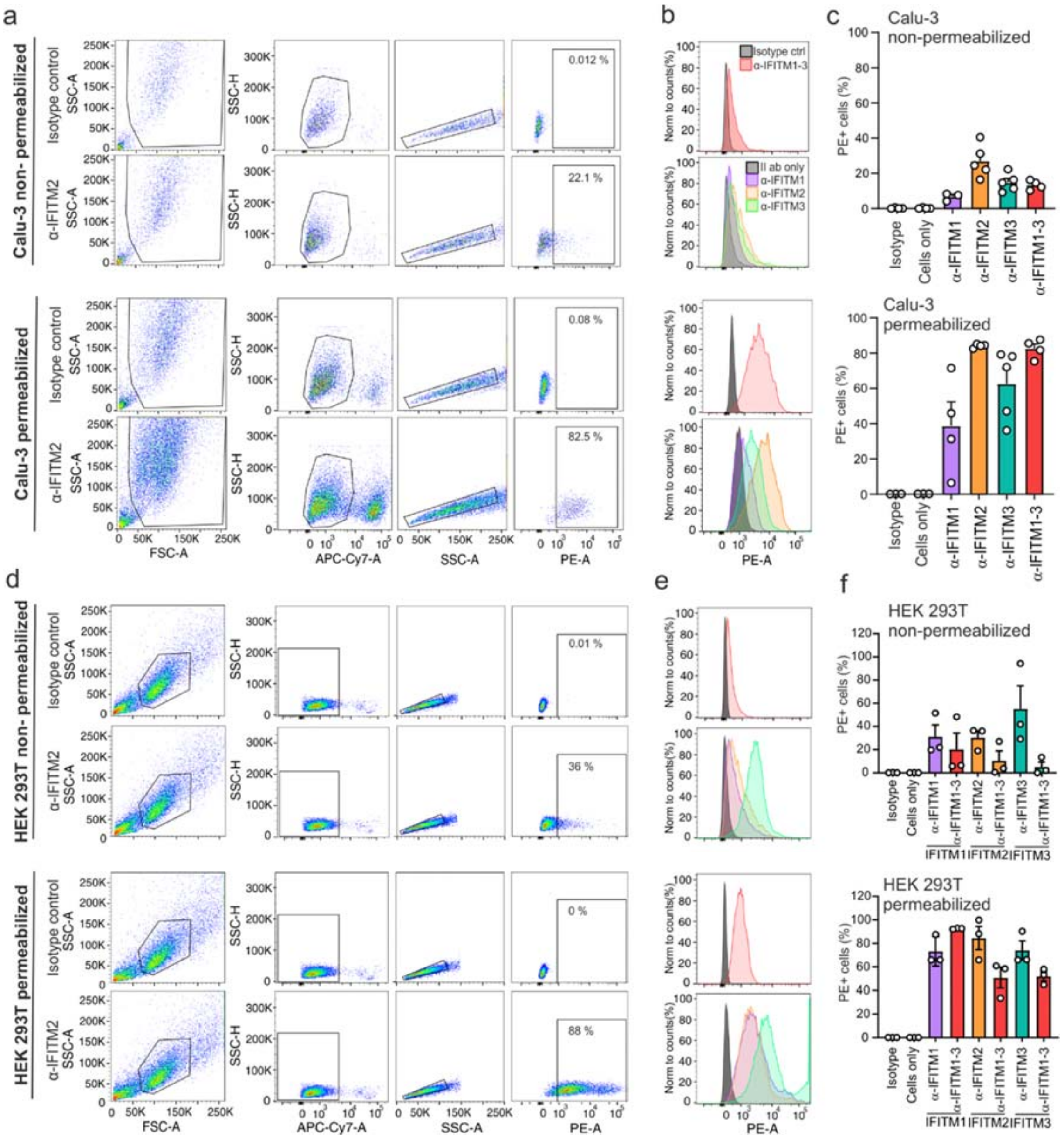


**Extended data Figure 7. Flow cytometric analysis of IFITM expression. a,** Gating strategy and flow cytometric detection of endogenous IFITMs in non-permeabilized (upper) or permeabilized (lower) Calu-3 cells using α-IFITM1, α-IFITM2, α-IFITM3 and α-IFITM1-3 antibodies. The left part shows examples for primary data and the bar diagrams quantitative results from the FACS analyses. **b and e,** Histograms represent the percentage of positive cells normalized for alive/single cells. **c,** Bars represent four independent experiments (±SEM) **d,** flow cytometric analyses of non-permeabilized (upper) or permeabilized (lower) HEK293T cells transiently transfected with constructs expressing the indicated IFITM proteins and analyzed using the corresponding α-IFITM1, α-IFITM2, α-IFITM3 or α-IFITM1-3, respectively. **f,** Bars represent three independent experiments (±SEM)

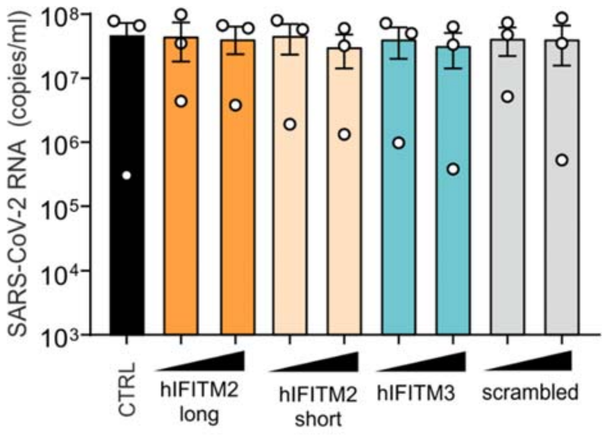


**Extended data Figure 8. Effect of IFITM-derived peptides on SARS-CoV-2 infectivity.** Viral N gene RNA levels in the supernatant of Calu-3 cells infected with SARS-CoV-2 pre-treated with two concentrations of IFITM-derived peptides. Bars represent two independent experiments measured in technical duplicates (mean, ±SEM).

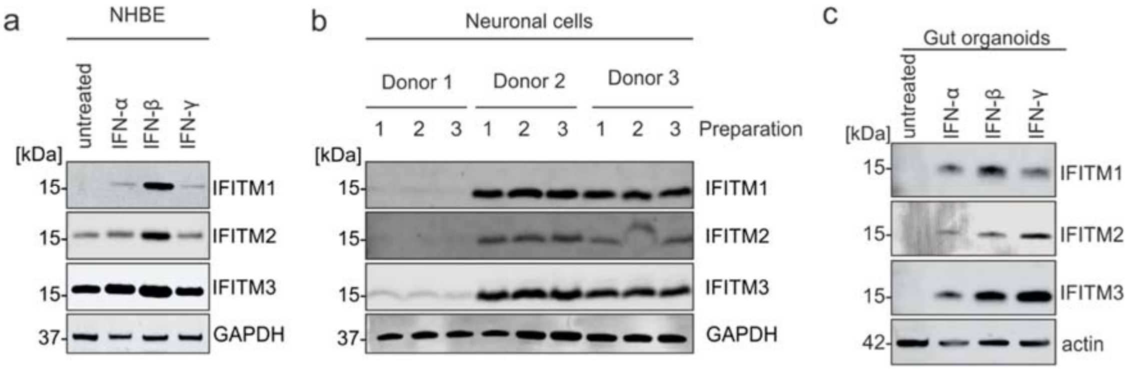


**Extended data Figure 9. Expression of IFITMs in primary lung cells, neuronal cells and gut organoids. a,** Expression of IFITM1, IFITM2 and IFITM3 in primary bronchial epithelial cells (NHBE) after stimulation with IFN-α2 (500 U/ml, 72 h), IFN-β (500 U/ml, 72 h) or IFN-γ (200 U/ml, 72 h). Immunoblot of whole cell lysates stained with anti-IFITM1, anti-IFITM2, anti-IFITM3 and anti-GAPDH. **b,** Expression of IFITM1, IFITM2 and IFITM3 in neuronal cells. Cell lysates have been prepared from three independent differentiations of induced pluripotent stem cells of three different donors. **c**, Expression of IFITM1, IFITM2 and IFITM3 after stimulation with IFN-α2 (500 U/ml, 72 h), IFN-β (500 U/ml, 72 h) or IFN-γ (200 U/ml, 72 h) in stem cell derived gut organoids.

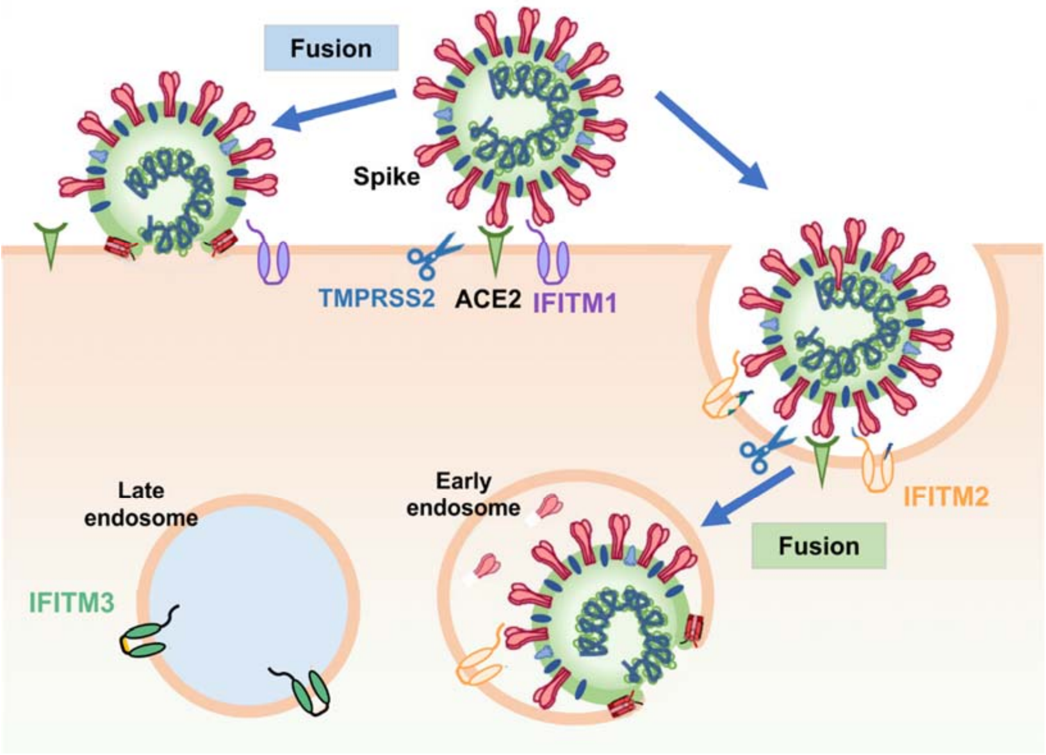


**Extended data Figure 10.** Schematic presentation of the potential role of IFITM protein in SARS CoV-2 infection. Modified from Ref. 24.
